# The APE2 nuclease is essential for DNA double strand break repair by microhomology-mediated end-joining

**DOI:** 10.1101/2022.07.21.500989

**Authors:** Hubert Fleury, Myles K. MacEachern, Clara M. Stiefel, Roopesh Anand, Colin Sempeck, Benjamin Nebenfuehr, Benjamin Dodd, Erin Taylor, Djelika Dansoko, Raquel Ortega, Justin W. Leung, Simon J. Boulton, Nausica Arnoult

## Abstract

Microhomology-mediated end-joining (MMEJ) is an intrinsically mutagenic pathway of DNA double strand break repair essential for proliferation of homologous recombination (HR) deficient tumors. While targeting MMEJ has emerged as a powerful strategy to eliminate HR-deficient (HRD) cancers, this is limited by an incomplete understanding of the mechanism and factors required for MMEJ repair. Here, we identify the APE2 nuclease as a novel MMEJ effector. We show that loss of APE2 blocks the fusion of deprotected telomeres by MMEJ and inhibits MMEJ in DNA repair reporter assays to levels comparable to Pol Theta suppression. Mechanistically, we demonstrate that APE2 possesses intrinsic flap-cleaving activity, that its MMEJ function in cells depends on its nuclease domain and further identify uncharacterized domains required for recruitment to damaged DNA. We conclude that HR-deficient cells are addicted to APE2 due to a previously unappreciated role in MMEJ, which could be exploited in the treatment of cancer.

## Introduction

Our DNA is subject to damage from endogenous sources, including reactive oxygen species or replication stress, or from external threats, such as UV light, environmental chemicals, or cosmic radiations. To cope with DNA damage and maintain genome stability, organisms have evolved an array of mechanisms, collectively termed the DNA damage response (DDR), that sense and repair DNA damage. Inaccurate DNA repair results in the accumulation of mutations or genomic rearrangement, which are early hallmarks of cancers. Furthermore, germline mutations affecting DNA repair pathways are associated with increased risk of developing different types of cancer (Knoch et al., 2012; O’Driscoll, 2012; Taylor et al., 2019). Similarly, acquired genetic or epigenetic alterations that compromise DDR promote cellular transformation and are found in a large proportion of cancers (Chae et al., 2016; Ma et al., 2018).

DNA double-strand breaks (DSB) are the most toxic type of lesion that occurs in DNA, and can be repaired by multiple pathways. Two-ended DSBs, which are created by radiations or nucleases, can be repaired by one of three DSB repair pathways, non-homologous end-joining (NHEJ), homologous recombination (HR) or microhomology-mediated end joining (MMEJ). MMEJ relies on the annealing of small microhomologies (3-20 bp) that flank the break site (Chang et al., 2017; Daley and Wilson, 2005) and that are exposed by short-range resection of the DSB, a step that is shared with HR (Truong et al., 2013). In mammals, MMEJ depends on the Polymerase Theta (Polθ, encoded by POLQ), which promotes annealing of the microhomologies and is responsible of the polymerization step (Ceccaldi et al., 2015; Chan et al., 2010; Kent et al., 2015; Mateos-Gomez et al., 2015; Mateos-Gomez et al., 2017; Wyatt et al., 2016). In addition to Polθ, PARP1, FEN1, XRCC1 and Ligase 3 have been implicated in MMEJ (Audebert et al., 2004; Liang et al., 2005; Sharma et al., 2015; Simsek et al., 2011; Wang et al., 2006). MMEJ is intrinsically error-prone, as microhomology annealing result in deletion of one of the two homologous sequences as well as the intervening DNA between the two microhomologies. Furthermore, Polθ is an inaccurate polymerase that frequently introduces mutations and can produce templated insertions (Carvajal-Garcia et al., 2020; Schimmel et al., 2017; Yu and McVey, 2010). MMEJ was first described as backup pathway because cells that are deficient in either NHEJ or HR become critically dependent on MMEJ for survival (Boulton and Jackson, 1996). However, low levels of MMEJ activity are detected at Cas9-induced DSBs in cells that have functional NHEJ and HR (Schep et al., 2021), and MMEJ repair scars are found in normal and cancer cells (Alexandrov et al., 2013; Ma et al., 2018; Schimmel et al., 2017).

While it is not fully understood why HR and MMEJ are synthetically lethal, a widespread hypothesis is the role of both pathways at single-ended DSBs (seDSBs). seDSBs occur when a replication fork converts an unrepaired single strand break and collapse. In unchallenged cells, they constitute the vast majority of unscheduled DSBs (So et al., 2017). Notably, repair of seDSBs by NHEJ is expected to induce aberrant chromosomal structures toxic to the cells, and NHEJ is indeed repressed at seDSBs (Britton et al., 2020; Chanut et al., 2016). The main repair pathway that is active at seDSBs is therefore HR (Britton et al., 2020), with MMEJ as a possible backup (Wang et al., 2019). Upon loss of HR, MMEJ likely becomes critical to repair these seDSBs, explaining the synthetic lethal interaction between HR and MMEJ. Supporting this model, MMEJ has been shown to repair seDSBs that are mediated by the Cas9^D10A^ or that arise from replication fork collapse (Wang et al., 2019).

HR deficiency results in high levels of genomic instability, which can in part be explained by the upregulation of the intrinsically mutagenic MMEJ (Abyzov et al., 2015; Malhotra et al., 2013; Mateos-Gomez et al., 2015; Schimmel et al., 2017; van Schendel et al., 2016; Zhang and Jasin, 2011). It is therefore not surprising that, among all the DNA repair genes that are frequently mutated in cancer, somatic mutations or epigenetic repression of HR genes are the most commonly found (Heeke et al., 2018). The most prominent example is high-grade serous epithelial ovarian cancers (HGS-EOC), in which HR is deficient in over 50% of tumors (Ledermann et al., 2016). Germline mutations in BRCA genes, PALB2 and Rad51 paralogs, all critical for the HR pathway, are also observed in ovarian, breast, prostate, and pancreatic cancers (Nguyen et al., 2020).

Seminal studies found that small molecule inhibitors of PARP confer selective killing of HRD tumors (Bryant et al., 2005; Farmer et al., 2005). This led to the clinical development of PARP inhibitors, which are now commonly used to treat patients with HRD HGS-EOC as well as metastatic breast, prostate, and pancreatic cancers (Siegel et al., 2020). However, only 50% of HRD tumors respond to PARPi treatment due to innate resistance and of those that respond most tumors eventually relapse due to acquired drug resistance (Li et al., 2020). Encouraging, POLQ inhibitors have been developed that could be used to treat PARPi resistant tumors, at least those that have lost the 53BP1-Shieldin pathway (Zatreanu et al., 2021). Because POLQ expression is very low or absent in normal cells but is over-expressed in many cancers, including those with HRD, it has emerged as a powerful therapeutic target complementary to PARP inhibitors (Higgins and Boulton, 2018). However, our understanding of MMEJ remains limited, and lacks a complete understanding of the factors that cooperate with Polθ to promote MMEJ repair. Filling this knowledge gap is critical, as the identification of unknown MMEJ proteins could provide alternative targets for treatment of HRD cancers.

Through genome-wide CRISPR-Cas9 screens (Doench et al., 2016) that exploit the synthetic lethal interaction between MMEJ and HR, we have identified the APE2 nuclease as a novel MMEJ effector. We show that loss of APE2 blocks the fusion of deprotected telomeres by MMEJ (Sfeir and de Lange, 2012) and inhibits MMEJ in DNA repair reporter assays to levels comparable to POLQ suppression.

## Results

To discover potentially uncharacterized effectors of MMEJ, we exploited the synthetic lethality observed when cells lose both HR and MMEJ (Ceccaldi et al., 2015; Mateos-Gomez et al., 2015). To identify genes that are synthetically lethal with HRD in general, rather than with one gene in particular, we compared the genetic interactions of two HR genes, *BRCA1* and *PALB2* (Venkitaraman, 2001; Xia et al., 2006; Zhang et al., 2009). We created isogenic cell lines by knocking out *BRCA1* or *PALB2* with transient CRISPR-Cas9 expression in HT1080 fibrosarcoma HR-proficient cells and isolated clones for each gene knockout (Figures S1A-S1C). As expected, the resulting HRD cells were sensitive to the PARP inhibitor Olaparib and failed to form Rad51 foci upon irradiation (Figures S1D-S1F).

We next performed genome-wide CRISPR-Cas9 dropout screens (Doench et al., 2016) in the HT1080, HT1080-*BRCA1*^KO^ and HT1080-*PALB2*^KO^ cells, which identified genes that are synthetically lethal with loss of *BRCA1* or with *PALB2* (Figures 1A-C and S2A-S2C). Plotting the depletion scores of both dropout screens against each other revealed those genes whose depletion conferred synthetic lethality in both HR-deficient lines (Figure 1D). Among the top candidates, we found *PARP1*, *XRCC1*, *NBN* (encoding Nbs1), *LIG3* and *POLQ* (encoding Polθ), which are already known to participate in MMEJ repair (Audebert et al., 2004; Ceccaldi et al., 2015; Dutta et al., 2017; Liang et al., 2005; Mateos-Gomez et al., 2015; Sfeir and Symington, 2015; Truong et al., 2013; Wang et al., 2006). FEN1, also suggested to be involved in MMEJ (Mengwasser et al., 2019; Sharma et al., 2015), is essential for Okazaki fragment processing and was lethal in the parental cells in our screening conditions, explaining why we did not find it as a significant hit.

**Figure 1:**
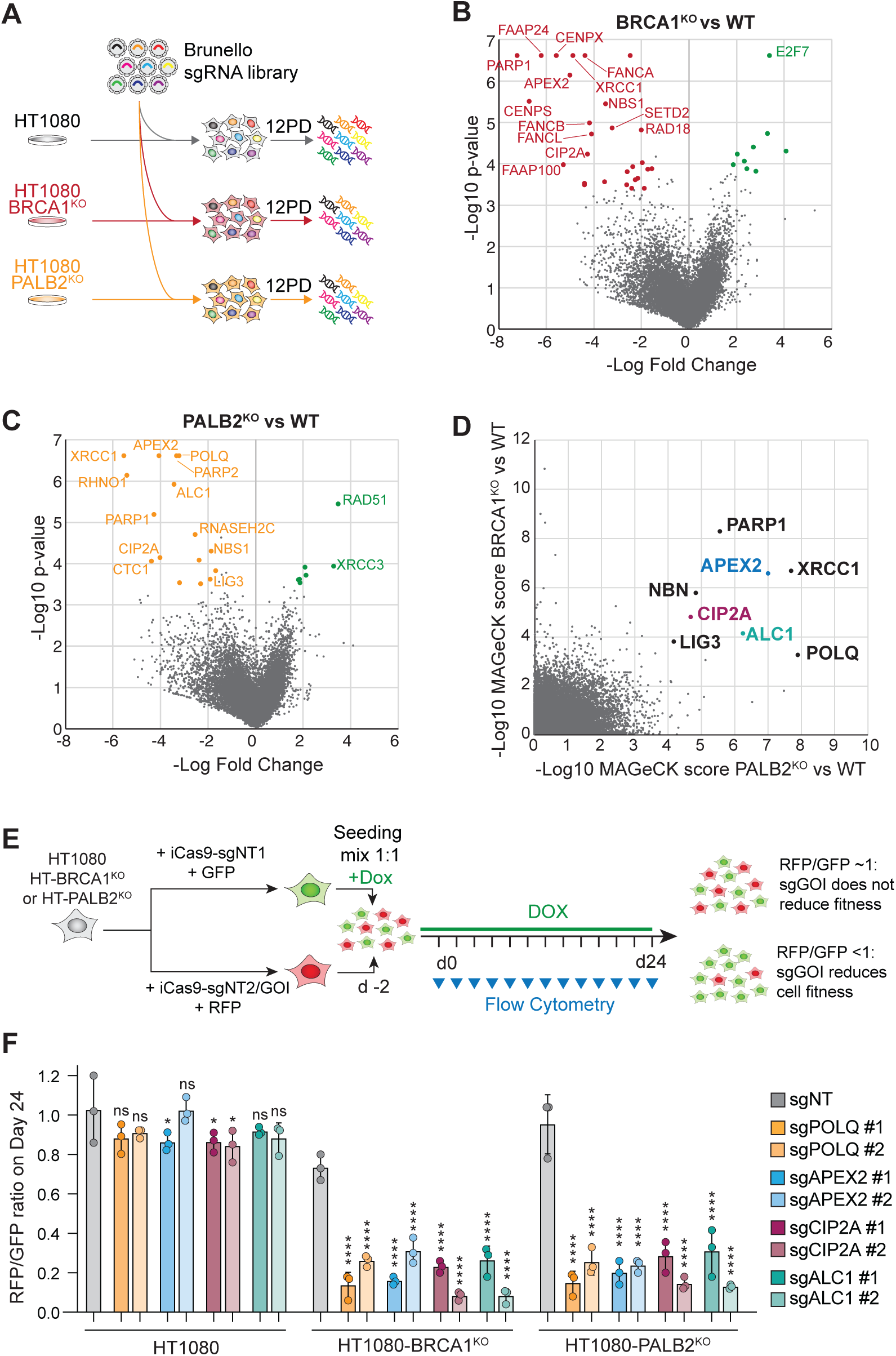
A dual CRISPR-Cas9 screens identifies MMEJ genes and APEX2, CIP2A and ALC1. **A.** Schematic of the dual CRISPR-Cas9 screen. **B.** Result of the genome wide CRISPR-Cas9 screen in HT1080-BRCA1^KO^ compared to parental HT1080. Most significant genes are annotated. **C.** Result of the genome wide CRISPR-Cas9 screen in HT1080-PALB2^KO^ compared to parental HT1080. Most significant genes are annotated. **D.** Result of the dual CRISPR-Cas9 screen: Plot of MAGeCK -Log10 depletion scores of *BRCA2*^KO^ vs *PALB2*^KO^. **E.** Schematic and timeline of the growth competition assay. HT1080, HT-*BRCA1*^KO^ or HT-*PALB2*^KO^ are transduced with iCas9, sgNT (non-targeting) and GFP or with RFP and the indicated sgRNA (sgGOI: gene of interest). GFP and RFP cells are mixed 1:1, treated with doxycycline (1µg/ml) to induce Cas9 expression, and subjected to flow cytometry every 2 days. **F.** Result of the growth competition assay at day 24. Values are % of RFP+ cells / % of GFP+ cells, normalized to day 0. 3 biological replicates. Data are mean ± s.d.

In addition to the previously reported MMEJ genes, we found *CIP2A*, *ALC1* and *APEX2* (encoding APE2) as significant hits (Figure 1D). The synthetic lethality between these three genes and HRD was previously reported by us and others (Adam et al., 2021; Alvarez-Quilon et al., 2020; Hewitt et al., 2020; Mengwasser et al., 2019; Verma et al., 2021). However, to our knowledge, their potential involvement in MMEJ has not been tested. CIP2A, in complex with TOPBP1, prevents mis-segregation of acentric chromosomes in HRD cells (Adam et al., 2021). ALC1 is a nucleosome remodeler that is essential during base excision repair (Ahel et al., 2009; Blessing et al., 2020; Hewitt et al., 2020; Verma et al., 2021). APE2 is a nuclease related to the apurinic/apyrimidic endonuclease APE1 and is involved in the repair of oxidative damage and at endogenous 3’ blocks that arise from removal of genomic ribonucleotides (Alvarez-Quilon et al., 2020; Burkovics et al., 2009; Burkovics et al., 2006; Hossain et al., 2018). Using two-color competitive growth assays (Noordermeer et al., 2018), clonogenic assays, and Annexin V/propidium Iodide staining, we confirmed that deletion of these three genes is lethal in *BRCA1*^KO^ and *PALB2* ^KO^ HT1080s but does not affect cellular fitness in the parental cells (Figures 1E, 1F, and S3). Next, we set out to determine whether CIP2A, ALC1 and/or APE2 also participate in MMEJ repair.

### APE2 is required for MMEJ-mediated fusions of deprotected telomeres

Due to the overwhelming competition with HR and NHEJ, accurately measuring MMEJ activity can be a challenge. To overcome this problem, we exploited fusion of telomeres as a readout of end-joining. Since the ends of linear chromosomes resemble DNA double strand breaks, cells have evolved the telomere-bound Shelterin complex to prevents the ends of chromosomes from activating DNA damage signaling and repair pathways (Arnoult and Karlseder, 2015). By selectively removing Shelterin, we can deprotect chromosome ends and activate DDR signaling and repair. For instance, TRF2 suppression leads to telomere fusions mediated by NHEJ, while MMEJ and HR remain inactive (Celli et al., 2006; van Steensel et al., 1998). Alternatively, MMEJ is redundantly suppressed by TRF1, TRF2 and the Ku heterodimer (Sfeir and de Lange, 2012). Their combined deletion specifically induces MMEJ-mediated fusions, while HR remains inhibited by RAP1 and POT1 (Palm et al., 2009; Sfeir and de Lange, 2012; Sfeir et al., 2010) and NHEJ is suppressed by the loss of Ku. This approach has been used to demonstrate Pol Theta’s central role in MMEJ (Mateos-Gomez et al., 2015). Here we exploited the deletion of TRF2 and Ku only, since it provides a reliably quantitative readout for MMEJ activity (Sfeir and de Lange, 2012).

The MMEJ-mediated telomere fusion assay must be performed in mouse cells, as deletion of the Ku heterodimer in human cells is lethal (Li et al., 2002). We therefore knocked out *XRCC5* (encoding Ku80) with CRISPR-Cas9 in conditional *TERF2^F/-^* Cre^ERT2^ MEFs (Celli and de Lange, 2005) and isolated a stable clone (Figure 2A). As expected, tamoxifen-induced deletion of *TERF2* in the parental MEFs led to DNA-PK-dependent telomere fusions, confirming that they are mediated by NHEJ. Conversely, when we induced TRF2 depletion in the *XRCC5*^KO^ MEFs, the telomere fusions were unaffected by DNA-PKcs inhibition, but suppressed upon treatment with the PARP inhibitor Olaparib, indicating that they are mediated by MMEJ (Figures 2A, 2B and Figures S4A and S4B). We employed this quantitative fusion assay to assess the potential contribution of APE2, CIP2A and ALC1 during MMEJ. Using inducible Cas9 and two different small guide RNAs (sgRNA) per gene, we knocked out *APEX2*, *CIP2A* and *ALC1* in *XRCC5*^KO^ *TERF2^F/-^* MEFs and induced telomere fusions with tamoxifen driven deletion of *TERF2* (Figures 2C-2E and Figure S4C). Treatment with Olaparib or *POLQ* knockout were used as positive controls of MMEJ inhibition. As expected, we found that *POLQ* knockout or PARP inhibition significantly reduced telomere fusions. Furthermore, Olaparib treatment in *POLQ* knockout cells did not have an additive effect. Deletion of CIP2A or ALC1 did not reduce MMEJ-mediated telomere fusions, indicating that these proteins are dispensable for MMEJ. In contrast, knockout of *APEX2* significantly repressed telomere fusions by MMEJ to the same extent as knockout of *POLQ.*, Furthermore, Olaparib treatment did not further reduce the fusions in sg*APEX2* cells. These data demonstrate that APE2 is as necessary as Polθ for MMEJ-mediated telomere fusions (Figures 2D and 2E). To determine whether APE2’s function in end-joining is specific to MMEJ, we performed the telomere fusion assay in the parental *XRCC5*^WT^ *TERF2^F/-^* MEFs. We found that NHEJ-mediated fusions were unaffected by knockout of *APEX2*, *CIP2A* or *ALC1* (Figure 2F). We conclude that loss of APE2 compromises end-joining by MMEJ but has no impact on NHEJ-dependent telomere fusion events.

**Figure 2.**
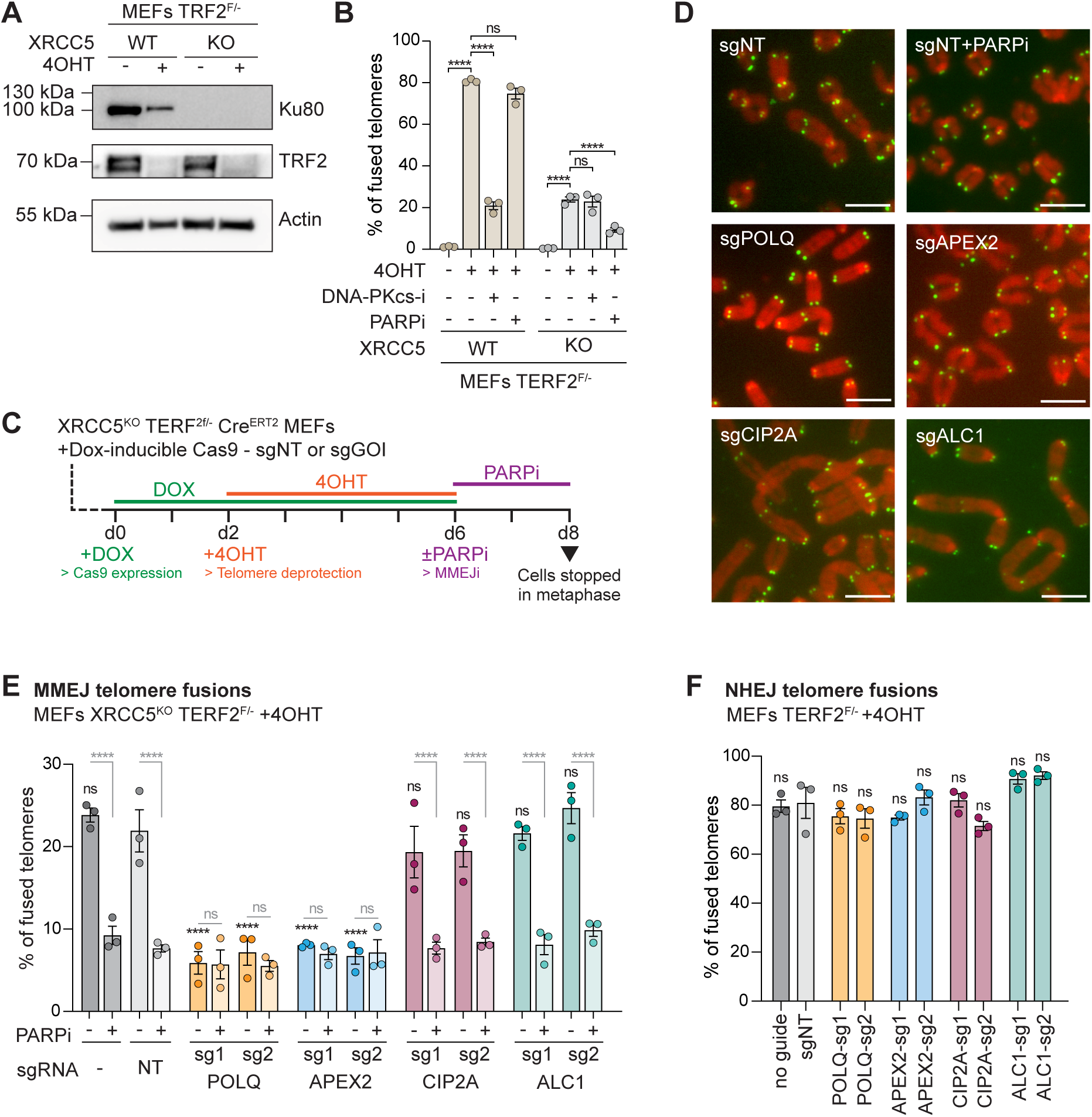
APE2 is required for MMEJ-mediated fusions of deprotected telomeres. **A.** Western Blot of Ku80, TRF2 and Actin in parental and *XRCC5*^KO^ *TERF2^F/-^* upon treatment with 4OHT (1µM). **B.** Percentage of fused telomeres in *XRCC5*^WT^ and *XRCC5*^KO^ *TERF2^F/-^* MEFs upon treatment with tamoxifen (4OHT, 1µM), DNA-PKcs-inhibitor (NU7441, 1µM) and PARP inhibitor (Olaparib, 20µM). 3 independent experiments. n=20 metaphases counted for each condition, each replica. Data are mean ± s.d. **C.** Experimental timeline for Figure 2D-E. sgNT: non-targeting small guide RNA. sgGOI: guide RNA targeting a gene of interest. **D.** Representative images of metaphase spreads analyzed in e. Red: DNA (DNA - DAPI), Green: telomere FISH. Scale bar: 5µm. **E.** Quantification of telomere fusions in *XRCC5*^KO^ *TERF2^F/-^*MEFs transduced with iCas9 and the indicated sgRNA, and treated with doxycycline (1µg/ml), 4OHT (1µM), and PARPi (Olaparib, 20µM). 3 independent experiments. n=22 metaphases counted for each condition, each replica. Data are mean ± s.e.m. Statistical analyses: Grey: untreated vs. PARPi. Black: Untreated sgNT vs. untreated no guide or sgTarget. **F.** Quantification of telomere fusions in *XRCC5*^WT^ *TERF2^F/-^* MEFs transduced with iCas9 and the indicated sgRNA, treated with doxycycline (1µg/ml) and 4OHT (1µM). 3 independent experiments. n=18 metaphases for each condition, each replica. Data are mean ± s.e.m. Statistical analysis for B, E and F: One-way ANOVA. ****p<0.0001.

### APE2 is a key effector of the MMEJ repair pathway

APE2’s essential function in MMEJ-mediated telomere fusions led us to explore a broader role of APE2 in MMEJ repair of intra-chromosomal DSBs. To test for such function, we first turned to fluorescent MMEJ DNA repair reporter assays. Using CRISPR-Cas9, we knocked out *POLQ* and *APEX2* in HT1080 cells, isolated three independent *POLQ*^KO^ and four *APEX2*^KO^ clones, and observed that their cell cycle remains unperturbed (Figures S5A-C). We created an MMEJ DNA repair reporter that consists of a lentiviral mCherry cassette interrupted by an ISCe1 site flanked by a 10 bp microhomology. Upon expression of ISce1, repair of the cut site by microhomology annealing leads to reconstitution of the mCherry cassette and fluorescence expression (Figure 3A and Figure S6A). We transduced this reporter into the knockout clones and established stable cell lines. We then transfected these cells with a vector expressing ISce1 and BFP, and measured the percentage of mCherry positive cells within the transfected cells by flow cytometry (by gating on BFP positive cells) (Figures 3B and 3C). We found that MMEJ was significantly reduced in all *APEX2*^KO^ clones when compared to the parental cells, and that the reduction in mCherry+ cells was comparable to that observed in *POLQ*^KO^ clones (Figure 3C).

**Figure 3.**
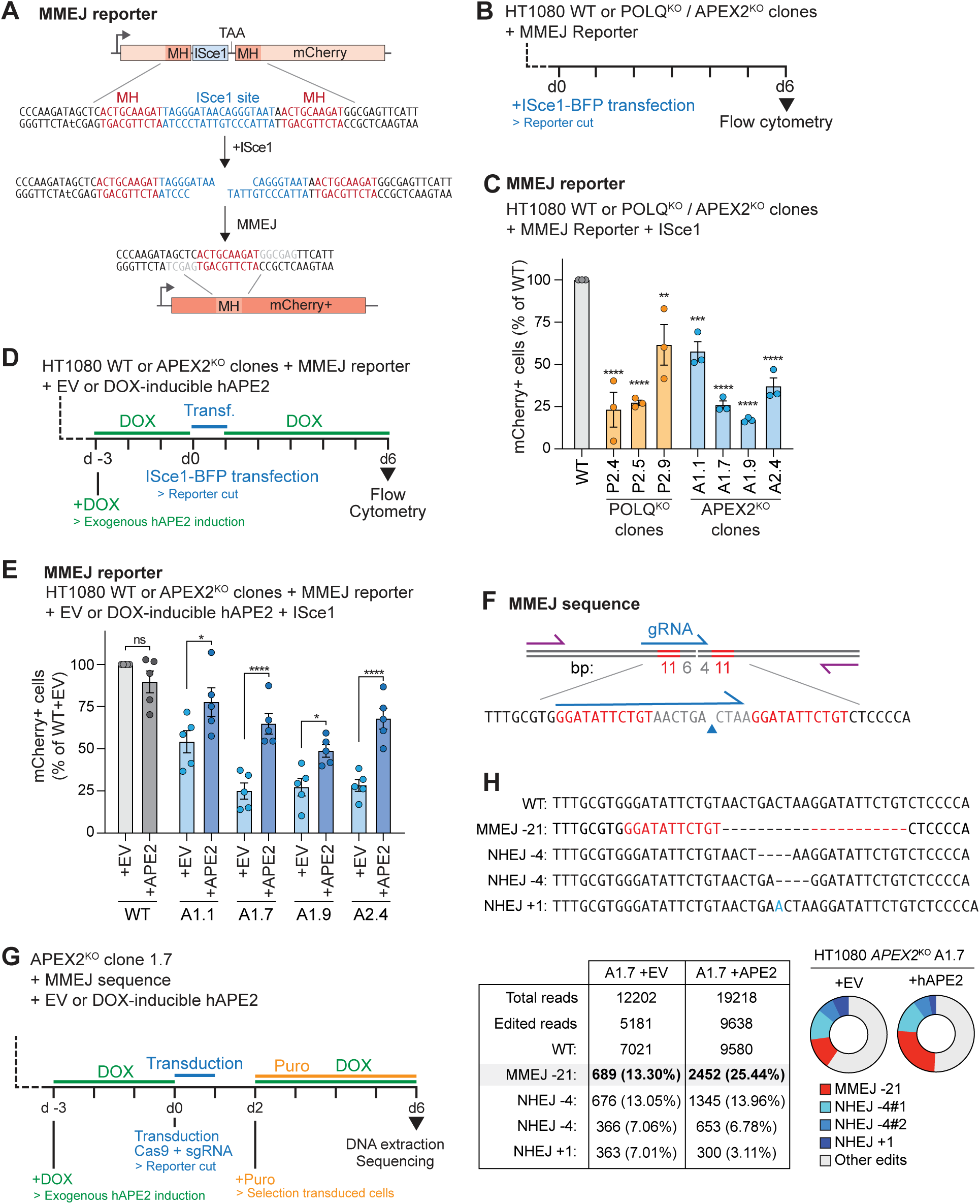
APE2 is a key effector of the MMEJ repair pathway. **A.** Schematic of the MMEJ repair reporter used in 3C, 3E, 5G, and 6E. Upon induction of a DSB by ISce1 and repair using annealing of the indicated microhomologies, the mCherry cassette is reconstituted. **B.** Experimental timeline for 3C. **C.** MMEJ quantification in parental HT1080 and indicated KO clones. mCherry+ cells (MMEJ+) were scored in the BFP+ population (I-Sce1+). Values are normalized to WT. 3 independent experiments. Data are mean ± s.e.m. **D.** Experimental timeline for 3E. **E.** MMEJ quantification in parental HT1080 and *APEX2*^KO^ clones complemented with empty vector (EV) or hAPE2. 5 independent experiments. Data are mean ± s.e.m. **F.** Schematic of the MMEJ sequence. **G.** Experimental timeline for 3H. **H.** Depiction and quantification of the WT and most frequent repair outcomes sequences in *APEX2*^KO^ clone A1.7 complemented with empty vector or hAPE2. Percentages represent % of edited total sequences. Statistical analysis for C and E: One-way ANOVA. *p<0.05, **p<0.01, ***p<0.001, ****p<0.0001.

To exclude potential off-target effects or clonal divergences, we complemented the parental cells and the *APEX2* knockout clones containing the MMEJ reporter with an empty vector (EV) or with exogenous APE2. While exogenous expression of APE2 had no effect on MMEJ activity in the parental cells compared to the cells transduced with an empty vector, re-expression of APE2 significantly increased the percentage of mCherry+ cells in all four *APEX2*^KO^ clones (Figures 3D and 3E). We next analyzed these isogenic cells using the EJ2 DNA repair reporter (Bennardo et al., 2008), which has been broadly used to measure MMEJ ((van de Kooij and van Attikum, 2021);Figure S6B). Again, we found that MMEJ activity is reduced in *POLQ* and *APEX2* knockout clones to similar extents, and that exogenous APE2 expression restored MMEJ in the *APEX2*^KO^ clones (Figures S6C-F). Finally, we designed a sequence containing an 11 bp microhomology spaced by 10 bp that can be cut by Cas9 and a guide RNA (Figure 3F) and transduced it into *APEX2*^KO^ cells complemented with empty vector or APE2. We then transduced these cells with Cas9-sgRNA, selected the transduced cells, and amplicon sequenced the repair junction (Figure 3G). We found that the percentage of edited reads corresponding to the MMEJ repair event (- 21bp deletion) were almost twice more frequent in cells complemented with APE2 (Figure 3H). Conversely, reads corresponding to the most frequent NHEJ repair outcomes, deletions of four nucleotides or addition of one A at the break site, were either found at similar frequency or reduced upon APE2 expression. Collectively, these data demonstrate that APE2 is a required for MMEJ repair of DSBs.

### APE2 possesses intrinsic 3’ flap-cleaving activity

The MMEJ pathway (Figure 4A) is initiated by short-patch resection of the DSB by the CtIP/MRN complex, resulting in the formation of 3’ overhangs (Truong et al., 2013). Polθ helicase has been proposed to remove RPA, thereby facilitating annealing of the microhomologies (Mateos-Gomez et al., 2017). Polθ polymerase activity then fills-in the gaps (Mateos-Gomez et al., 2015), and finally XRCC1/LIG3 complete ligation (Audebert et al., 2004). FEN1 is proposed to remove the 5’ flaps created by Polθ polymerization (Liang et al., 2005; Sharma et al., 2015). When micro-homologies do not directly flank the break, their annealing results in the formation of 3’ flaps that must be removed in order for Polθ to proceed to polymerization. In yeast, these 3’ flaps are removed by Rad1-Rad10, the orthologs of human XPF-ERRC1 (Ma et al., 2003). XPF-XRCC1 was shown to remove 3’ flaps during NHEJ-independent class switch recombination in B cells (Bai et al., 2021). However, its depletion has only minor effects on MMEJ activity at intrachromosomal breaks in mammals (Bennardo et al., 2008), suggesting that another nuclease is involved. Because APE2 is a 3’ to 5’ exonuclease with endonuclease activity, we reasoned that it could participate in the removal of the 3’ flaps that form upon microhomology annealing. However, a biochemical activity of hAPE2 on this type of structure has not been previously demonstrated. To test whether APE2 could remove flaps, we expressed and purified human wild type APE2 and the catalytic dead mutant APE2^D197N^ (Erzberger et al., 1998; Hadi et al., 2002) from insect cells (Figure 4B). Both 3’ flap and Y structure are predicted DNA intermediates during MMEJ repair and need to be removed for MMEJ completion (Figure 4A). Therefore, we tested the *in vitro* activity of APE2 on 3’ flap and Y-structure substrates, both containing 24 nt ssDNA region. Strikingly, we found that APE2 is capable of cleaving both types of structures *in vitro*, suggesting that it could function in MMEJ during flap removal (Figure 4C).

**Figure 4:**
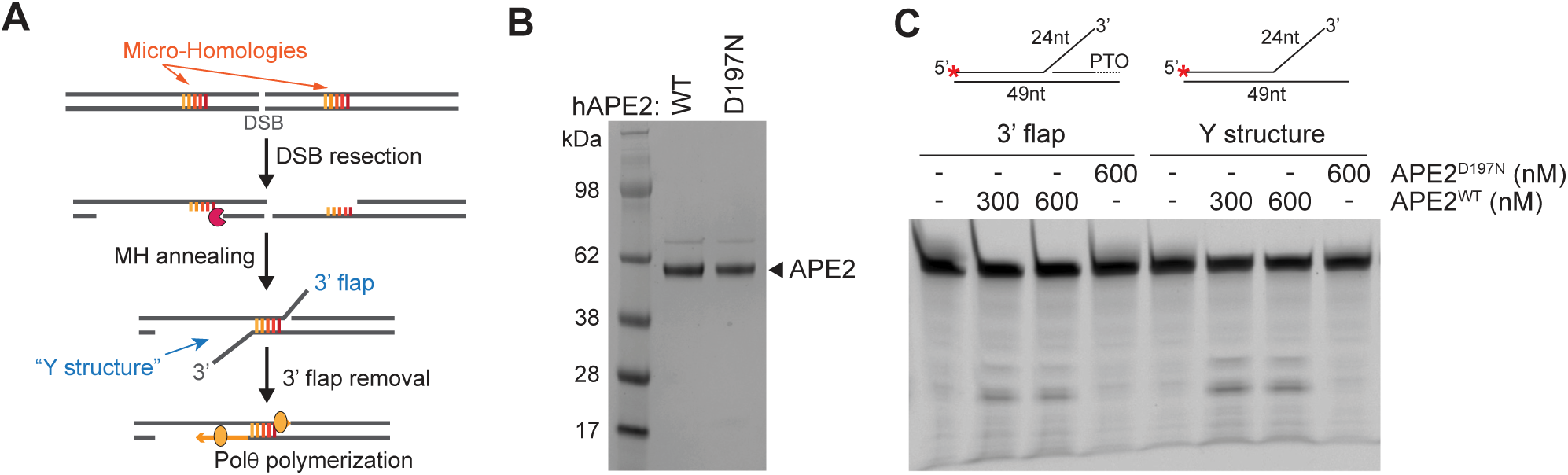
Human APE2 possesses intrinsic 3’ flap cleavage activity. **A.** Schematic of the MMEJ repair pathway. Depending on the length of resected DNA relative to microhomologies, 3’ flaps or Y structures (the strand opposite the flap is single stranded) can form upon microhomology annealing. **B.** SDS-PAGE gel (4-12% polyacrylamide) showing purified hAPE2 WT and hAPE2 D197N from insect cells. The gel was stained with Coomassie brilliant blue (CBB). **C.** Representative gel (15% polyacrylamide denaturing urea gel) of the 3’ flap and Y structure cleavage by APE2. The asterisk indicates the position of the fluorescein isothiocyanate (FITC) labelling at the 5’ end of the oligo. PTO indicates the presence of 4 phosphorothioate bonds at the 3’ end of complementary short oligo in 3’ flap. PTO bonds makes DNA refractory to nuclease cleavage.

### The PIP and Zinc-finger motifs of APE2 are dispensable for MMEJ

APE2 belongs to the ExoIII family of nucleases (Hadi and Wilson, 2000). Its catalytic domain, which possesses endonuclease, 3’-5’ exonuclease, and phosphodiesterase activities (EEP), is related to the base excision repair (BER) AP endonuclease APE1. However, APE2 displays stronger exonuclease and weaker AP endonuclease activities (Hadi et al., 2002; Unk et al., 2000; Unk et al., 2001). Beyond its catalytic domain, APE2 also possesses a C-terminal domain that is absent in APE1, which contains a PCNA interacting peptide (PIP) box and a zinc-finger motif (Zf-GRF). The interaction of APE2 with PCNA through its PIP box facilitates its recruitment to sites of oxidated DNA damage (Burkovics et al., 2009). In vitro, APE2-PCNA interaction promotes the phosphodiesterase and exonuclease activities of APE2 at 3’ resected DNA (Burkovics et al., 2009; Unk et al., 2002; Wallace et al., 2017). The APE2-PCNA interaction is also necessary for the removal of 3’ blocks arising during removal of genomic rNTPs *in vitro* (Alvarez-Quilon et al., 2020). The Zinc-finger motif promotes the recruitment of APE2 to ssDNA, as well as the subsequent PCNA-mediated exonuclease activity of APE2. Conversely, mutations in either the PIP box or the Zf-GRF domain do not affect APE2’s endonuclease activity at abasic sites (Burkovics et al., 2009; Wallace et al., 2017). To determine if the PIP box and/or Zf-GRF domains are required for MMEJ repair, we generated a series of mouse and human APE2 mutants. We introduced two mutations, D197N (CD1) and D277A (CD2), that suppress all three catalytic activities of APE2 (Alvarez-Quilon et al., 2020; Burkovics et al., 2006; Hadi et al., 2002), we deleted the PIP box (ΔPIP) and the zinc-finger motif (ΔZf-GRF), or introduced a mutation in the Zing-finger (Zf-GRF^mut^) that abolishes its ssDNA binding capabilities (Wallace et al., 2017) (Figure 5A). First, we stably complemented the *XRCC5*^KO^ *TERF2^F/-^* iCas9-sg*APEX2* MEFs with empty vector (EV) or with Doxycycline-inducible wild-type or mutant mAPE2 (Figure 5B). Treatment with DOX enables simultaneousinduction of Cas9 expression (APE2 knockout) and exogenous expression of APE2. We then induced telomere deprotection by deleting TRF2 with tamoxifen and quantified the percentage of MMEJ-mediated telomere fusions (Figures 5C and 5D). As expected, we observed very few telomere fusions in APE2 knockout cells without complementation (∼5% fused telomeres), while exogenous APE2 expression restored the levels of fusions to that observed in wildtype cells (∼20%). Expression of either catalytic dead APE2 mutants could not rescue the levels of fusions, indicating that the catalytic activity of APE2 is critical for its role in MMEJ. Conversely, we observed that expression APE2ΔPIP fully rescued telomere fusions, showing that the PIP box is dispensable for MMEJ-mediated fusions. The zinc-finger deletion or mutation displayed an intermediate phenotype, suggesting that MMEJ-driven repair is only partially dependent on a functional Zf-GRF. To determine whether the PIP and the zinc-finger motifs could have a redundant role in the recruitment of APE2 to DSBs and its MMEJ function, we also complemented cells with an exogenous mAPE2 that carried both the PIP deletion and the zinc-finger mutation (ΔPIP-Zf^mut^) (Figures 5A and 5B). We found that deletion of the PIP motif had no additional effect on telomere fusions compared to the zinc-finger mutant alone, suggesting that the APE2-PCNA interaction is not required for MMEJ repair (Figure 5D). To extend these observations to intra-chromosomal break repair, we complemented HT1080-*APEX2*^KO^ cells containing the MMEJ reporter with wildtype or mutant human APE2 (Figure 5E), transfected the cells with ISce1 and measured MMEJ activity by flow cytometry (Figure 5F). Using this method, we obtained similar results as with the telomere fusion assay (Figure 5G). Collectively, these data show that MMEJ repair requires APE2 nuclease activity but not its interaction with PCNA.

**Figure 5.**
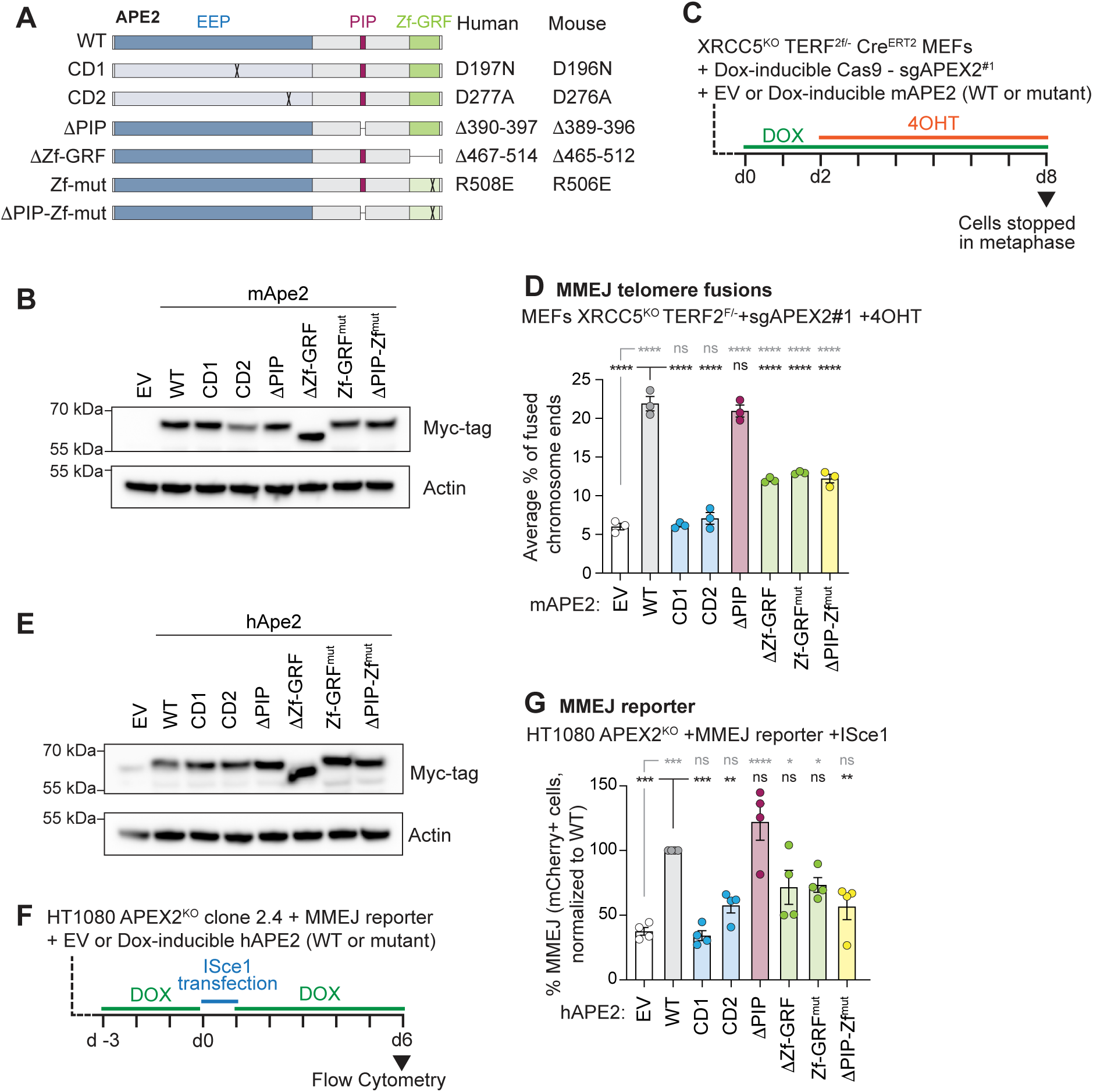
The APE2-PCNA interaction in dispensable for MMEJ. **A.** Schematic of wild type and mutants APE2. Amino acids mutated or deleted are indicated for both human and mouse APE2. **B.** Western Blot of Myc-tag (mAPE2) and Actin for the indicated MEFs used in D. **C.** Experimental timeline for Figures 5D and 6D: On day 0, Doxycycline treatment induces expression of Cas9 (*APEX2* knockout) and mAPE2 exogenous expression. On day 3, addition of 4OHT induces *TERF2* Cre recombination and telomere deprotection. On Day 8, cells are briefly treated with Colcemid and metaphase spreads are prepared. **D.** Quantification of telomere fusion in *XRCC5*^KO^ *TERF2^F/-^* MEFs transduced with iCas9, sg*APEX2*#1, and the indicated mAPE2 construct, and treated with doxycycline (1µg/ml) and 4OHT (1µM). EV: empty vector. 3 independent experiments. 26 metaphases were counted in each condition, each replica. Data are mean ± s.e.m. **E.** Western Blot of Myc-tag (hAPE2) and Actin for the indicated HT1080s used in G. **F.** Experimental timeline for Figures 5G and 6E. Cells were treated 3 days with Doxycycline to induce expression of Cas9 (*APEX2* knockout) and of the hAPE2 exogenous constructs prior to transfection with ISce1-BFP. 24 hours post-transfection, media was replaced (with Dox). Flow cytometry was performed 6 days after ISce1 transfection. **G.** MMEJ quantification using reporter from 2a in HT1080-*APEX2*^KO^ clone A2.4 complemented with indicated hAPE2. mCherry+ cells (MMEJ+) were scored in the BFP+ population (I-Sce1+). Values are normalized to WT. 4 independent experiments. Data are mean ± s.e.m. Statistical analysis for D and G: One-way ANOVA. Grey: vs empty vector, Black: vs WT. *p<0.05, **p<0.01, ***p<0.001, ****p<0.0001.

### Two uncharacterized conserved regions of APE2 mediate its recruitment to sites of damage

Since the PIP box and zinc-finger motif of APE2 are largely dispensable for MMEJ activity, we searched for other domains of APE2 that could mediate its recruitment to sites of DNA damage. First, we analyzed APE2’s sequence for conserved regions and discovered that, beyond the already known EEP, PIP and Zf-GRF domains, two other amino acid stretches showed a high level of conservation (Figure 6A) and are predicted structural domains by AlphaFold2 (Jumper et al., 2021) (Figure 6B). We therefore tested whether these conserved regions, which we named CR1 and CR2, are involved in MMEJ. We created CR1 and CR2 deletion mutants and tested their capacity to complement *APEX2* knockout in telomere fusion assays and DNA reporter assays (Figures 6C-E). We found that both CR1 and CR2 are required for MMEJ activity at deprotected telomeres and intra-chromosomal DSBs. We then examined whether these mutants impair the recruitment of APE2 to sites of damage. We fused GFP to WT, ΔCR1-, or ΔCR2- hAPE2, transfected U2OS cells with these constructs, and monitored their recruitment to sites of 405 nm laser-induced micro-irradiation upon BrdU treatment, a technique that induces DNA breaks, more frequently double stranded. First, we found that wild-type APE2 is efficiently recruited and rapidly accumulates to micro-irradiation sites (Figures 6F, S7A and S7B). Noticeably, deletion of the CR2 domain weakened APE2’s recruitment, while deletion of CR1 completely abolished it (Figures 6F and S7B). APE2^ΔCR1^ showed a more diffuse localization, suggesting that CR1 may facilitate the nuclear localization of APE2. To exclude the possibility that impaired recruitment to damage is due to a compromised nuclear import, we fused the ΔCR1 mutant to a nuclear localization domain and found APE2^ΔCR1^-NLS still showed a diminished recruitment to sites of damage (Figures 6F and S7B). We then performed the reciprocal experiment and tested whether CR1 alone is sufficient to trigger recruitment to sites of damage. We fused GFP to a small amino acid patch containing CR1 (amino-acids 313-352) and found that this short protein domain was able to recruit GFP to sites of damage as well as full length APE2 (Figures 6G and S7C). Collectively, these data show that recruitment of APE2 to DSBs requires CR1 and to a lesser extent CR2 and are necessary for MMEJ.

**Figure 6.**
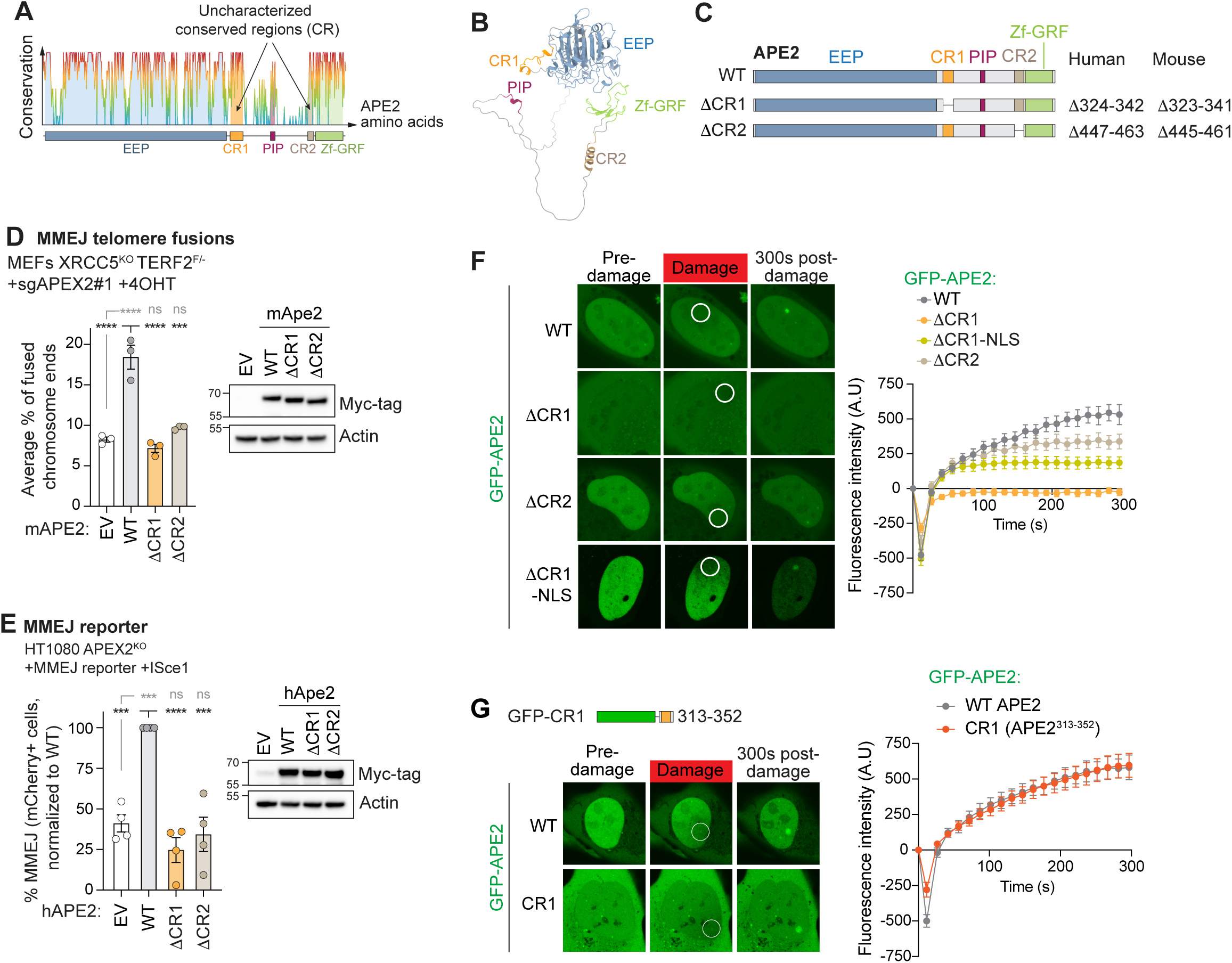
Two conserved regions of APE2 promote its recruitment to damage sites and repair by MMEJ. **A.** Schematic of hAPE2 amino acid conservation. Two uncharacterized conserved regions are indicated. **B.** AlphaFold2 structure prediction of mAPE2. **C.** Schematic of APE2 CR1 and CR2 deletion mutants. Deleted amino acids are indicated for both human and mouse APE2. **D.** Quantification of telomere fusion in *XRCC5*^KO^ *TERF2^F/-^*MEFs transduced with iCas9, sg*APEX2*#1, and the indicated mAPE2 construct, and treated with doxycycline (1µg/ml) and 4OHT (1µM). EV = empty vector. 3 independent experiments. Data are mean ± s.e.m. Right: Western blot showing expression of the mAPE2 constructs. **E.** MMEJ quantification using the MMEJ repair reporter in HT1080-*APEX2*^KO^ clone A2.4 complemented with indicated hAPE2. mCherry+ cells (MMEJ+) were scored in the BFP+ population (I-Sce1+). Values are normalized to WT. 4 independent experiments. Data are mean ± s.e.m. Right: Western blot showing expression of the mAPE2 constructs. **F-G.** Recruitment of GFP-hAPE2 to sites of laser micro-irradiation (405 nm, 60% power) upon treatment with BrdU (10µM, 20h) in U2OS cells transfected with the indicated constructs. Data show mean of all cells analyzed over 3 independent experiments. In F, N=26 (WT, 1′CR1-NLS), 24 (1′CR1), or 29 (1′CR2) cells. In G, N=21 (WT) or 20 (CR1) cells. Statistical analysis for D and E: One-way ANOVA. Grey: vs empty vector, Black: vs WT. ***p<0.001, ****p<0.0001.

### APE2 motifs participation in MMEJ predicts their requirement for HRD cell survival

The synthetic lethal interaction between loss of APE2 and HRD was previously assumed to be due to APE2’s function in base excision repair or ribonucleotide excision repair (RER). In light of its core function in MMEJ, and the known synthetic lethal interaction between MMEJ and HR, we sought to determine whether the dependence of HRD cell on APE2 is actually due to its MMEJ repair function. We showed above that the PIP box of APE2 is dispensable for MMEJ activity. Conversely, the APE2-PCNA interaction has been previously shown to be required for the BER and RER activities of APE2 (Alvarez-Quilon et al., 2020; Burkovics et al., 2009; Willis et al., 2013 {Unk, 2002 #24767). Furthermore, MMEJ is partially active with the zinc-finger mutants, while the corresponding mutation abrogates Xenopus APE2’s nucleolytic activity on 3’ resected DNA as well as APE2-mediated ATR/Chk1 activation upon oxidative stress (Wallace et al., 2017; Willis et al., 2013). We therefore tested whether the domains that are dispensable for MMEJ activity but necessary for BER and RER are similarly necessary or dispensable for survival in HRD cells. We performed a fluorescent growth competition assay in *BRCA1*^KO^ and *PALB2*^KO^ cells after complementation of sg*APEX2* with the different hAPE2 mutants (Figure 7A-C). We found that the catalytic activity of APE2 as well as the two CR domains are essential for survival of HRD cells. The zinc-finger mutants showed an intermediate phenotype, similarly to what we observed for MMEJ activity. Importantly, we found that APE2ΔPIP could fully rescue cellular survival, and that the double ΔPIP-Zf^mut^ mutant displayed a similar impact on cellular fitness as the zinc-finger mutation alone. We conclude from these results that a domain’s importance in APE2’s MMEJ activity correlates with its requirement for HR-deficient cells survival. These results strongly suggest that HRD cells are critically dependent on APE2 because of its role in MMEJ, rather than due to its function in other DNA repair pathways.

**Figure 7.**
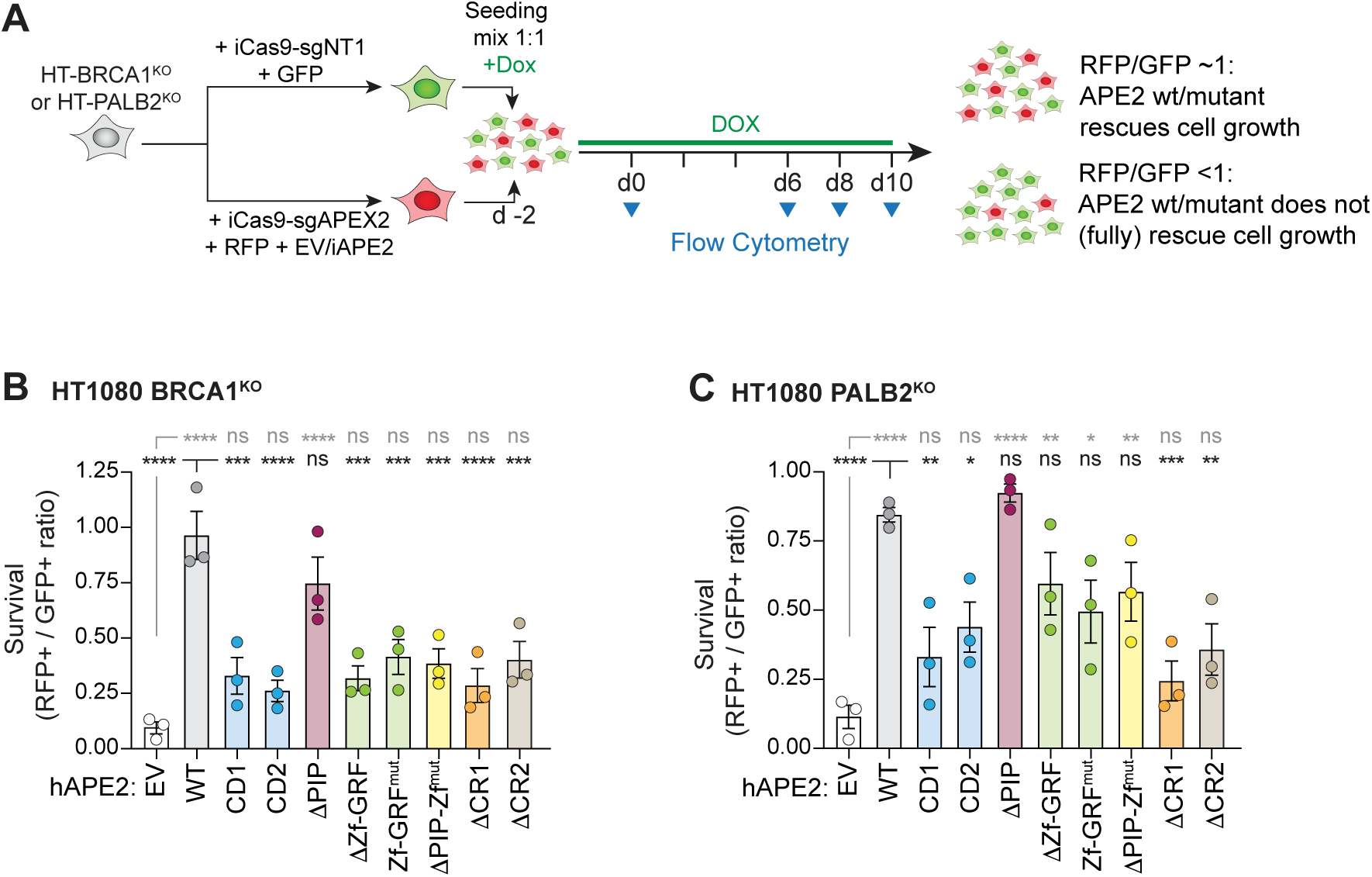
APE2 domains’ requirement for MMEJ predicts their necessity for survival in HR- deficient cells. **A.** Schematic and experimental timeline of the growth competition assay. HRD cells are transduced with sgNT (non-targeting) and GFP or with sg*APEX2*#1, RFP and the indicated hAPE2 or empty vector (EV). **B-C.** Growth competition assay in HT1080-*BRCA1*^KO^ cells (B) and HT1080-*PALB2*^KO^ cells (C). Values are % of RFP+ cells / % of GFP+ cells, averaged on days 6, 8 and 10 and normalized to day 0. 3 independent experiments. Data are mean ± s.e.m. Statistical analysis: One-way ANOVA. Grey: vs empty vector, Black: vs WT. *p<0.05, **p<0.01, ***p<0.001, ****p<0.0001.

## Discussion

We demonstrate here that the nuclease activity of APE2 is essential for the repair of DSBs by MMEJ, establishing APE2 as a novel and critical player for the MMEJ driven repair mechanism. We show that APE2 possesses intrinsic flap cleavage activity (Figure 4), suggesting that 3’ flap removal could be its function in vivo. Other nucleases, especially XPF-ERCC1, have been proposed to perform the step of 3’ flap removal during MMEJ. Deletion of Rad1-Rad10, the yeast ortholog of ERCC1-XPF1, greatly reduces MMEJ repair, suggesting a role for this nuclease complex in 3’ flap removal. It was also recently shown that the 3’ flap activity of XPF-ERCC1 is required for class switch recombination (CSR) in NHEJ-deficient mouse B cells (Bai et al., 2021). However, ERCC1-XPF is not required for MMEJ repair of intrachromosomal DSBs (Bennardo et al., 2008). An alternative possibility is that ERCC1-XPF functions in Polθ-independent alternative end-joining. Supporting this hypothesis, earlier studies found that CSR levels are not affected by suppression of Polθ, XRCC1, or Ligase III in B cells with inactivated NHEJ (Boboila et al., 2012a; Boboila et al., 2012b; Yousefzadeh et al., 2014), suggesting that NHEJ-independent CSR may mainly rely on Polθ-independent repair.

In addition, APE2 could play a role in the DSB resection. Although CtIP/MRN is known to promote the initial resection of DSBs (Truong et al., 2013), one could imagine that APE2 is required during G1, when CtIP is not robustly phosphorylated by CDKs (Ira et al., 2004; Yun and Hiom, 2009). Indeed, while CtIP can be phosphorylated in G1 by PL3K, this phosphorylation seems to promote resection-dependent NHEJ rather than MMEJ (Barton et al., 2014; Biehs et al., 2017). Several lines of evidence refute this hypothesis. First, we show here that APE2 is required for MMEJ-mediated fusion of deprotected telomeres (Figure 2). Since telomeres naturally bear a long 3’ overhang (Makarov et al., 1997; McElligott and Wellinger, 1997), resection is unlikely to be a limiting step for the initiation of MMEJ at telomeres. Secondly, in contradiction to a potential requirement of APE2 specifically in G1, recent evidence suggest that DSBs induced during interphase are repaired by MMEJ in mitosis, upon PLK1-mediated CtIP phosphorylation (Llorens-Agost et al., 2021; Wang et al., 2018). Finally, while the APE2-PCNA interaction is necessary for processive exonuclease activity of APE2 (Burkovics et al., 2009; Unk et al., 2002; Wallace et al., 2017), we show here that the APE2-PCNA interaction is fully dispensable for MMEJ, both at deprotected telomeres and at intrachromosomal DSBs (Figure 5).

Using DNA damage recruitment as a functional readout, we observed that APE2, like Polθ (Mateos- Gomez et al., 2015), is swiftly accrued to laser-induced micro-irradiation damage sites (Figure 6). Using the same assay, we show that two conserved structural modules, CR1 and CR2, are required for the recruitment of APE2 to sites of DNA damage. Notably, CR1 promotes the translocation of APE2 from cytoplasm to nucleus, which plays a substantial role in promoting APE2 damaged chromatin accumulation. CR1 and CR2 are respectively 18 and 16 amino acids. Therefore, the most likely mechanism by which they promote the recruitment of APE2 to DNA is through protein-protein interactions, though a direct DNA interaction cannot be fully excluded.

Finally, we show that the PIP box of APE2 is dispensable for MMEJ and survival of HRD cells (Figures 5 and 7). On the contrary, the interaction with PCNA stimulate the *in vitro* 3’-5’ exonuclease and phosphodiesterase activities of yeast Apn2 and human APE2 (Burkovics et al., 2009; Unk et al., 2002). Likewise, upon H2O2-mediated oxidative stress, the PIP box aids APE2’s recruitment to sites of damage and is required for APE2-mediated activation of ATR, ATRIP and Chk1 (Burkovics et al., 2009; Willis et al., 2013). Regarding 3’ blocking lesions, while APE2 alone can remove Top1cc upon proteolysis degradation, removal of intact Top1cc and the final 3’ resection step both require the presence of PCNA (Alvarez-Quilon et al., 2020). Finally, during ribonucleotide excision repair, APE2 either directly removes 2’-3’ cyclic phosphate formed after the first TOP1 cleavage, or it removes the DNA-protein crosslink left after a second TOP1 cleavage. APE2’s activity on both types of substrates is very limited in the absence of PCNA (Alvarez-Quilon et al., 2020). In summary, the PCNA-APE2 interaction had been involved in almost all the previously known APE2 repair activities. Yet, we and others (Alvarez-Quilon et al., 2020) have found that the PIP box of APE2 is fully dispensable for survival of HR deficient cells, suggesting that another APE2 function, independent of PCNA, was driving its synthetical lethality with HR. To date, MMEJ is the only known repair activity of APE2 that does not require interaction with PCNA. Consistently, we show that an APE2 lacking the zinc-finger motif provides partial rescue to both MMEJ activity, as well as partial survival of HR-deficient cells (This study and (Alvarez-Quilon et al., 2020)). These data suggest that the synthetic lethality between APE2 and HR is due to APE2’s function in MMEJ, rather than its role in other repair mechanisms. Conversely, the synthetic lethality between APE2 and RNAseH2 is likely mediated by APE2’s function in ribonucleotide excision repair (Alvarez-Quilon et al., 2020). The in vitro processing of these lesions by APE2 depends on the presence of PCNA, suggesting that the APE2- PCNA interaction is likely required for survival of RNAseH2 deficient cells. Further work should test whether APE2ΔPIP can rescue cellular fitness in APE2/RNAseH2 DKO. Likewise, APE2 has been implicated in the removal of TOP1-mediated 3’ blocks, a function that is partially redundant with TDP1 (Alvarez-Quilon et al., 2020). It would be similarly useful to ascertain whether APE2’s PIP box is necessary to prevent sensitivity to camptothecin upon TDP1 suppression.

In conclusion, considering that HR-deficient cells are fully dependent on MMEJ and that our data point at APE2 as core component of the MMEJ pathway, APE2 represents a potential therapeutic target to eliminate HRD cancers, either as an alternative to or in combination with POLQ or PARP inhibitors (Zatreanu et al., 2021).

## Limitations of the study

We have focused here on the function of APE2 during MMEJ at double-ended DSBs. However, the lethality of the APE2-HR double knockout occurs without exogenous DNA damage, suggesting a role of APE2 during replication-coupled repair. Likewise, Polθ-mediated MMEJ was shown to repair DSBs caused by replication fork collapse (Wang et al., 2019). Therefore, future studies should evaluate the contribution of APE2 and its MMEJ function during replication stress. Furthermore, we have identified two novel domains that mediate APE2’s recruitment to sites of laser-induced micro-irradiation damage and established their essential function in MMEJ. It will be interesting, in the future, to test whether these domains also play a role in the recruitment of APE2 to other types of DNA damage and repair, such as oxidative damage and ribonucleotide excision repair.

## STAR METHODS

### KEY RESOURCES TABLE

See Supplementary materials

### RESOURCE AVAILABILITY

#### Lead contact

Further information and requests for resources and reagents should be directed to and will be fulfilled by the lead contact: Nausica Arnoult

#### Materials availability

Materials used in this study are available upon reasonable request.

#### Data and code availability

Any additional information required to reanalyze the data reported in this paper is available from the lead contact upon request.

### EXPERIMENTAL MODEL AND SUBJECT DETAILS

#### Cell lines, cell culture and drugs

HT1080 were derived from patient diagnosed with fibrosarcoma and purchased from ATCC (CCL-121). MEFs TRF2F/- Rosa26-CreERT2 were established in the de Lange lab (REF Celli 2005) and obtained from ATCC (CRL-3317). Lenti-X 293T were derived from a transformed human embryonic kidney cell line and purchased from Takara (#632180). All these cell lines and their derivates were maintained in a low oxygen condition (3% O2 and 7.5% CO2) and grown in DMEM medium (Corning # 10-013CV) with 10% Bovine Calf Serum (Seradigm #2100-500),100 μg/mL Penicilin/Streptomycin (Lonza #09-757F) and 1X MEM Nonessentiel Amino Acids (Corning#25-025CI). U2OS were derived from a moderately differentiated sarcoma, purchased from ATCC (HTB-96), and cultured in Dulbecco’s modified Eagle’s medium (DMEM) at 37°C with 5% CO2.

Drugs used are: Doxycycline (1µg/ml, TCI #D4116), Olaparib (20µM, Selleckchem, AZD2281 #S1060), DNA-PKi (1µM, NU7441, R&D Systems, #3712/10), 4-Hydroxy-tamoxifen (1µM,4OHT, Cayman chemical company #17308)

### METHOD DETAILS

#### Plasmids

For CRISPR-Cas9 mediated knocks-out, except when stated otherwise, sgRNAs were cloned into pLenti-Guide-iCas9-Blast, a modified form of TLCV2(Barger et al., 2019) (A gift from Adam Karpf, Addgene #87360) in which eGFP was removed and puromycin was replaced by blasticidin resistance. Guides cloning was done following the golden gate cloning protocol and verified by Sanger sequencing. For APE2 exogenous expression, Myc-tagged codon-optimized human and mouse APEX2 were obtained by DNA synthesis (gBlocks, IDT DNA), and cloned using InFusion (Takara) into pLenti-rtta-Hygro, a modified form of pLIX_403 (A gift from David Root, Addgene #41395) in which Puro was replaced by Hygromycin selection and the bases 547-2297 were replaced by the myc-tagged constructs. All the different mutant versions of APE2 were then obtained by PCR and InFusion cloning.

To create the MMEJ reporter (pLenti-Puro-MMEJ_rep), we used a modified form of pLenti-puro(Guan et al., 2011) (A gift from le-Ming Shih, Addgene #39481) in which the bases 2525-2982 were replaced by CMV enhancer/promoter - MMEJ reporter – WPRE. The MMEJ reporter is made of mCherry- T2A-BFP in which the mCherry sequence is interrupted by with an I-SceI restriction site flanked by 10 bp of microhomology. The sequence was obtained by DNA synthesis (IDT DNA), and cloning was performed using InFusion (Takara).

The MMEJ Sce1 reporter was further modified to create pLenti-Neo-MMEJ_seq, in which we replaced the puromycin selection by neomycin, and replaced the microhomology-ISce1 site by 10 bp microhomologies flanking 11 bp that can be cut with Cas9. For cloning we used synthetic DNA gBlocks from IDT DNA and InFusion (Takara).

All plasmids were verified by restriction digest and Sanger sequencing.

#### Lentiviral transduction and infection

To produce lentivirus, 500,000 Lenti-X 293T cells were in one well of a 6-well plate. The next day, cells were transfected with packaging plasmids (0.5µg pCMV-VSV-G, 1.1µg pD8.9), 0.85µg transfer plasmid, 7.35µg of PEI MAX (Polysciences Inc. - Cat #: 24765-1) and 10mM HEPES. Media was refreshed 24h later with 3ml of complete DMEM. Virus-containing supernatant was collected 48h and 72h post transfection, cleared through a 0.45µm PES Filter membrane (Whatman Uniflo #9914-2504) and directly used to infect cells in the presence of 5 µg.ml polybrene (EMD Millipore #TR-1003-G). Forty-eight hours after infection, cells were washed and selected with 1 μg/ml Puromycin (Alfa Aesar #J61278- MC), 500 μg/ml G418 (neomycinR) (Cornung #30-234-CR), 200μg/ml Hygromycin B (Biosciences #31282-04-9), or 10 μg/ml Blasticidin (RPI #B12200-0.05). Selection was performed until complete death of control uninfected cells.

#### Knock-out cells

For knock-out of BRCA1 and PALB2, HT1080 cells were subjected to three rounds of transfection with pX330(Cong et al., 2013) (A gift from Feng Zhang, Addgene #42230) containing sgBRCA1 or sgPALB2, using Lipofectamine 2000 (ThermoFisher Scientific #11668027). Individual clones were isolated and pre-screened on sensitivity to Olaparib and by Western blot. Finally, selected clones were verified by allele sequencing: Total genomic DNA was extracted using 30µl of Quick extract (Biosearch Technologies – Cat #: QE09050) and PCR amplified around the Cas9 cut region using KOD master mix (EMD Millipore Corporation – Cat #: 71842-3). The PCR products were cloned into pCR™Blunt II- TOPO™ using the Zero Blunt™ TOPO™ PCR Cloning Kit (ThermoFisher #450245), transformed into DH5 competent cells and at least 12 bacterial clones were analyzed by Sanger sequencing.

For knock-out of XRCC5, MEFs TRF2F/- Rosa26 CreERT2 were transduced with pLentiGuide- iCas9-Puro, a modified form of TLCV2 (Barger et al., 2019) (A gift from Adam Karpf, Addgene #87360) in which eGFP was removed, and the XRCC5 gRNA was cloned. Individual cellular clones were isolated and screened by western blot.

For knock-out of APEX2 and POLQ, HT1080 cells were transduced with pLentiGuide-iCas9-Blast containing a guide RNA targeting APEX2 or POLQ, individual cellular clones were isolated and analyzed by PCR amplification and TOPO cloning followed by sanger sequencing.

#### CRISPR-Cas9 screens

For each isogenic cell line (Parental HT1080, HT1080-BRCA1^KO^, and HT1080-PALB2^KO^), 150 million cells were spinfected (Joung et al., 2017) for 2 hours at 1,000g and 33°C in 24-well plates (3.2 million cells per well in 3 ml of medium and 10 µg/mL of polybrene) with the Brunello genome-wide CRISPR/Cas9 library(Doench et al., 2016) (viral particles were directly obtained from Addgene, #179- LV73) at an MOI of 0.3. 24 hours later, cells were trypsinized and transferred to 15 cm tissue culture petri dishes, then selected the next day with 0.6µg/mL puromycin. Selected cells were then regularly split and counted, with minimal seeding of 45 million cells, and cultured for 12 population doublings after selection. Sequencing and analysis: Genomic DNA were then isolated using QIAamp DNA Blood Kits (QIAGEN #51104), sonicated, guide RNA sequences were captured using the Fli-Seq Library prep kit (Eclipse Bio), and PCR amplified with indexes and adapters for Illumina sequencing. The library was sequenced on an Illumina NovaSeq through Novogene Co. Paired-end reads were aligned used Bowtie 2 (Langmead and Salzberg, 2012), and the gRNA counts were counted using a lab developed Python script. Count files were then analyzed using MAGeCK (Li et al., 2014).

#### Telomeric fusion assay

Six days after treatment with 4-hydroxy-tamoxifen, MEFs were synchronized in metaphase for 3 hours with 0.1 µg/ml colcemid (Gibco KaryoMAX, #15212-012), media and cells were collected and pooled, centrifuged and subjected to hypotonic choc in 10ml 75mM KCl for 30 minutes at 37°C, then fixed and washed 3 times in methanol::glacial acetic acid 3::1 (v/v). Metaphases were then dropped onto superfrost microscope slides and dried overnight. For FISH, slides were rehydrated 10 min in PBS, fixed with 3.7% formaldehyde for 2min, washed 5 minutes in PBS, then treated 10 minutes at 37°C with 1 mg/ml pepsin in citric acid pH2, washed 3 times in PBS and fixed again in formaldehyde for 2 min. After three PBS washes, the slides were dehydrated in consecutive ethanol baths (75%, 95% and 100%, 3 minutes each) and allowed to air dry. Slides were then layered with 40 μl of 0.3 ng μl−1 Alexa488-OO- (CCCTAA)3 PNA probe (PNA Bio Inc. #F1001), diluted in 70% (v/v) deionized formamide; 0.25% (v/v) blocking reagent (NEN); 10 mM Tris pH 7.5; 4.1 mM Na2HPO4; 450 μM citric acid; 1.25 mM MgCl2), denatured for 4 min at 76 °C and incubated for 2 h at room temperature. Slides were then washed twice for 15 min in 70% (v/v) formamide; 10 mM TrisHCl pH 7.5 and three times for 5 min in 50 mM TrisHCl pH 7.5; 150 mM NaCl; 0.08% Tween-20. Slides were mounted with Vectashield Plus Dapi (Vector Laboratories H-1900).

#### MMEJ Reporter assay

Parental HT1080, POLQ^KO^, and APEX2^KO^ clones were transduced with pLenti-Puro-MMEJ_rep and selected with puromycin. Parental HT1080s and APEX2^KO^ clones containing the MMEJ reporter were then transduced with pLenti-rtta-Hygro-EV or pLenti-rtta-Hygro-hApe2, or mutant APE2 constructs (Clone A1.7 only). Cells were then transfected with pCVL.Sce-T2A-BFP (A gift from Andrew Scharenberg obtained from Addgene #32627) using lipofectamine 3000 (following manufacturer’s instructions). Reporter cells were split and passaged for 6 days, then analyzed by FACS for BFP and mCherry expression. The percent of mCherry+ and BFP+ cells was quantified from gated BFP+ cells that received the I-SceI virus.

#### MMEJ sequence and MiSeq-Sequencing

Clones A1.7 were transduced with pLenti-Neo-MMEJ_seq, selected with G418, then complemented with the pLenti-rtta-Hygro-EV or pLenti-rtta-Hygro-hApe2 and selected with hygromycin. Doxycycline was used to induced hApe2 expression. Three days after dox treatment, cells were transduced with pLenti-CRISPRv2-Puro (Stringer et al., 2019) (a gift from Brett Stringer, Addgene #98290) in which the corresponding guide RNA to cut the MMEJ sequence was cloned. Cells selected with puromycin were then harvested and total genomic DNA was extracted using Quick extract (Biosearch Technologies – Cat #: QE09050). For the first PCR, we used primers amplifying a sequence of 78bp around the Cas9 cut region of the reporter. The first PCR was performed in 25µl reactions, each consisting of 2x KOD master mix (EMD Millipore Corporation – Cat #: 71842-3), 300 ng of template DNA, and 0.25 μM of primers for 30 cycles. PCR purifications were performed using AMPure XP beads (Beckman Coulter Inc – Cat #: A63881) at a ratio of x2. For the second PCR, we diluted the amplicons from the first PCR 20-fold and used these amplicons as a template for the primers that are attached to the Illumina MiSeq adapter sequences and an 8-bp barcode sequence as an index to distinguish each PCR amplicon. The second PCR was performed in 25µl reactions, each consisting of 2x KOD master mix, 1.25 µl of PCR1 and 0.25 μM of primers during 10 cycles. PCR purifications were then performed using AMPure XP beads at a ratio of x1.4. Concentrations were calculated by Qubit dsDNA High Sensitivity Kit (Invitrogen – Cat #: Q32854) and the different amplicons were mixed to obtain a 4nM pooled library solution. The quality and size were analyzed by 2100 BioAnalyzer High Sensitivity DNA kit (Agilent Technologies – Cat #: 5067- 4626).

Pooled libraries at 4nM were denatured and diluted to 8pM with 20% PhiX control (Illumina – Cat #: FC-110-3001) per the Illumina Denature and Dilute Libraries Guide Protocol A (Illumina Document # 15039740v10). 8pM denatured library and MiSeq Reagent Kit (Illumina – Cat #: MS-103-1002) were loaded into the MiSeq system (Illumina – Cat #: SY-410-1003) per the Illumina MiSeq System Guide (Illumina Document # 15027617v06). Sequencing data was paired-end aligned using the program PEAR (Zhang et al., 2014) with the following options: -j 8. Paired-end aligned reads were then trimmed using the program cutadapt (Martin et al., 2006) with the following options: -j 8 -e 2 -g ^GCTAAGTTGAAGGTGACGAAGGG…CGTTACCCAAGATTCAAGCCTGCAAGAT$ -m 40 -M 110 – discard-untrimmed. Trimmed reads were then sorted, counted, and analyzed using a lab developed Python script.

#### Growth competition assay

Indicated sgRNAs were cloned into LentiGuide-iCas9-Blast-eGFP or LentiGuide-iCas9-Blast-RFP, a modified form of TLCV2 (REF: Barger Cancers 2019) (addegene #87360) in which Cas9-T2A-eGFP and EF1a-Puro were replaced by Cas and EF1a-Blast-T2A-RFP or EF1a- Blast -T2A-eGFP. After lentiviral infection and blasticidin selection, mCherry and GFP-expressing cells were mixed 1:1 (15,000 cells each) and seeded in a 12-well plate with doxycycline (1µg/ml) to induce Cas9 expression. Cells were then harvested for flow cytometry analysis of GFP and mCherry fluorescence on the indicated day, with d0 corresonding to 48 hours post Cas9 induction.

#### Biochemical assays

To prepare construct for the expression of recombinant APEX2, coding region of APEX2 with fused N-terminal 8Xhis tag and C-terminal Twin-StrepII tag was synthesized by and purchased from GeneWiz. The fused tags were flanked by PreScission protease site to remove them if needed. The construct was codon optimized for the expression in *Spodoptera frugiperda* (Sf9) insect cells. The synthesized region was inserted into pACEBac1 vector using BamHI and XbaI restriction sites. To prepare APEX2 variant D197N, QuickChange site directed mutagenesis kit was used according to manufactures instructions (Agilent). The introduction of desired mutation in plasmids were confirmed via DNA sequencing.

To express APEX2 in *Spodoptera frugiperda* (Sf9) cells, bacmid and baculoviruses were prepared according to manufactures recommendation (Bac-to-bac system, Life Technologies). For protein expression, 800 ml cells were seeded at 500,000/ml and infected with APEX2 baculovirus next day at 1x106 cells/ml after 24 hrs. The infected cells were further incubated at 27° C for 52 hrs with continuous agitation. To collect cell pellet, cells were spun at 500 g for 10 min, washed once with 1xPBS (137 mM NaCl, 2.7 mM KCl, 10 mM Na2HPO4, 1.8 mM KH2PO4). The pellet was snap frozen and stored at -80° C. To purify the protein, the harvested cell pellet was thawed on ice. All subsequent steps were carried out either 4° C or in cold room. The thawed pellet was resuspended with 3 volume of lysis buffer (50 mM Tris-HCl pH 7.5, 1 mM ethylenediaminetetraacetic acid (EDTA), protease inhibitor cocktail tablets (Roche), 30 µg/ml leupeptin (Merck), 1 mM phenylmethylsulfonyl fluoride (PMSF), 1 mM dithiothreitol (DTT), 20 mM Imidazole) in cold beaker and incubated for 15 min with continuous stirring. To resuspended cells, 16.6% glycerol and 310 mM NaCl were added to final concentration (in this order). The cell suspension was further incubated for 30 min for lysis with continuous stirring. Next, cell suspension was centrifuged at ∼ 48000 g for 30 min to obtain soluble extract. In the meantime, Ni-NTA (nickel-nitrilotriacetic acid) resin (Qiagen) was pre-equilibrated with Ni-NTA wash buffer (50 mM Tris-HCl pH 7.5, 1 mM β-ME, 300 mM NaCl, 1 mM PMSF, 10% glycerol, and 20 mM Imidazole). The pre-equilibrated Ni-NTA resin was added to soluble extract in 50 ml falcon tubes and incubated for 1 h with continuous rotation. The resin was washed 4x batchwise with centrifugation at 2000 g for 2 min with Ni-NTA wash buffer. The resin was transferred to chromatography columns (Biorad) and washed twice on column with Ni-NTA wash buffer. The protein was eluted from resin with Ni-NTA elution buffer (Ni-NTA wash buffer containing 150 mM Imidazole). The elution was further diluted with 4 volumes of dilution buffer (50 mM Tris-HCl pH 7.5, 1 mM β-ME, 300 mM NaCl, 1 mM PMSF, 10% glycerol). The diluted sample was incubated with pre-equilibrated Strep-tactin XT 4Flow resin (IBA) for 1 h with continuous mixing. Post incubation, strep resin was transferred to chromatography column and washed 6x with strep wash buffer (50 mM Tris-HCl pH 7.5, 1 mM β-ME, 300 mM NaCl, 1 mM PMSF, 10% glycerol). Finally, protein was eluted with strep elution buffer (50 mM Tris-HCl pH 8.0, 1 mM β-ME, 300 mM NaCl, 1 mM PMSF, 10% glycerol, 50 mM biotin) and dialyzed overnight in dialysis buffer (50 mM Tris-HCl pH 7.5, 1 mM β-ME, 150 mM NaCl, 1 mM PMSF, 10% glycerol). Next day, dialyzed sample was aliquoted, snap frozen and stored at -80 C until further use. All APEX2 variants were simultaneously purified with WT protein using identical protocol.

The nuclease assays were carried out in 15 µl buffer containing 40 mM Tris-HCl pH 8.0, 1 mM MgCl2, 1 mM MnCl2 1 mM DTT, 0.1 mg/ml bovine serum albumin (BSA, New England Biolabs) and 5’ end FITC (Fluorescein isothiocyanate) labelled 25 nM DNA substrate (in molecules). During all steps, samples were protected from light as much as possible. The reactions were assembled on ice and proteins were added, mixed, and incubated at 37° C for 30 mins. The reactions were stopped by adding 1 ul Proteinase K (Roche, 18.4 mg ml-1) and 0.5 uL of 0.5 M EDTA in each sample, and incubating sample for 30 mins at 37° C. Finally, 15 uL loading buffer (95% formamide, 20 mM EDTA, and bromophenol blue) was added to all samples and products were separated on 15% polyacrylamide (19:1 acrylamide-bisacrylamide, Bio-Rad) denaturing urea gels. The gels were directly imaged in ChemiDoc MP imaging system.

All DNA oligonucleotides were commercially synthesized and purchased from Merck Life Sciences. To prepare all substrates, DNA oligoes were mixed in 1:1 ratio in 1x annealing buffer (50 mM Tris-HCl pH 7.5, 100 mM NaCl, 5 mM MgCl^2+^) boiled at 95° C for 3 min and gradually cooled down overnight to room temperature. The names and sequences of DNA oligoes used in this study is as follow: Oligo 1 (5′ FITC-AGCTACCATGCCTGC ACGAATTAAGCAATTCGTA ATCATGGTCATAGCT), Oligo 32_PTO (CGGTACCGATGGGTAAAG*T*A*G*G), Oligo 24_ROX (CCTACTTTACCCATCGGTACCGTGCTTAATTCGTGCAGGCATGGTAGCT-ROX-3’). (*) denotes phosphorothioate bonds in oligo and ROX indicates the presence and position of rhodamine. However, ROX label was not used during imaging of denaturing gels. The combination to prepare various substrates is as follows; 3’ flap (oligo 1+ oligo 24_ROX + oligo 32_PTO), Y structure (oligo 1 + oligo 24_ROX).

#### Laser micro-irradiation

APEX2 cDNA was subcloned into Gateway compatible destination vectors using Gateway cloning technology (Invitrogen, 11789100, 11791020). Mutations were created using Q5 Site-Directed Mutagenesis (New England Biolabs, R0176L, M0201S, M0202L, B0202S) according to the manufacturer’s instructions and verified using Sanger sequencing. U2OS cells were seeded into 6-well plates and transfected with GFP-APEX2 constructs. Then, 24 h after transfection, cells were moved to glass-bottom dishes (Cellvis) and pre-sensitized with 10 μM 5-bromo-2’-deoxyuridine (BrdU) in normal media 20 h prior to experimentation. Cells were damaged by laser-induced microirradiation using a Nikon Ti2 inverted fluorescent microscope and C2+ confocal system, as previously described (West et al., 2019). Live cell images were captured in 15-sec intervals. The fluorescence intensity of GFP-tagged protein in the damaged region was measured and normalized to the undamaged area in the same cell. Quantification analyses were done using NIS Elements Advanced Research software (Nikon). Analyses will be performed in GraphPad Prism.

#### Cellular growth assays

For IncuCyte cell proliferation phase-contrast imaging assay, HT1080s were seeded in 96-well plates at 1,000 cells per well. 24 hours later, cells were incubated with different concentrations Olaparib and imaged every 6 hours for 84 hours by phase contrast using the IncuCyte™ Live-Cell Imaging System (IncuCyte HD). Images were taken in four separate regions per well using a 10X objective.

For clonogenic assays, cells were seeded at 1,000 cells/well in 6-well plates and treated 24 hours later with different concentrations of Olaparib or with 1µg/ml of doxycycline, and kept in drugs for 8 days. Cells were then fixed with cold methanol and colored with a mix of 50% v/v methanol and 0.5% m/v brilliant Blue G (Sigma–Aldrich #B0770-SG), and colonies were counted under a stereomicroscope.

#### RT-qPCR

Total RNA was extracted from harvested MEFs using Trizol (Thermofisher Scientific #15596026), according to manufacturer’s protocol. One microgram of total RNA was subjected to reverse transcription using the Luna Script RT SuperMix (New England Biolabs #M30102). One microliter of the reverse transcribed product was diluted (1:10) and subjected to Q-PCR using sequence specific primers (400 nM) and Luna Universal QPCR MasterMix (New England Biolabs #M3003S). qPCR was performed using PCR Applied Biosystems - 7500 Fast Real-Time PCR with 50 °C/2 min, 95 °C/2 min, 40 cycles at 95 °C/15 s, 60 °C/1 min and 72 °C/30s). Gene expression values were normalized to GAPDH. Three independent experiments were performed in duplicate.

#### Flow for cell cycle analysis

Cultured cells were treated for 10 min with 30 μM BrdU (AlfaAesar #H27260-06), trypsinized, centrifuged and resuspend in 500µl PBS. Drop by drop 1.5ml ice-cold 100% ethanol was added to the cells. Pepsin (Sigma #p7000) (20min at 37degree) and HCL (20 min at room temperature) treatment were performed before labelling with Rat-anti-BrdU (AbD serotec #OBT0030S), Goat anti-rat-FITC (Southern Biotech #3030-02) and propidium iodide (AlfaAesar #J667664-MC). Flow cytometry was performed using MACSquant Analyzer 10 (Miltenyi) and analysed with FlowJo10.

#### Immunoblotting

Whole cell lysates were prepared by scraping cells with cold PBS. Protein extracts were done by using RIPA containing a protease and phosphatase inhibitor cocktail (Roche_05892791001). Protein concentration was measured using the bicinchoninic acid (BCA) protein assay (Thermo Fisher Scientific #23227). Thirty micrograms of total protein extracts were separated on Bolt 4-12% Bis-Tris Plus gels (Invitrogen #NW04120) and transferred onto PVDF membrane (Amersham Hybond P 0.2µm PVDF, GE Healthcare Life science #10600057). Membranes were blocked with 5% skim milk in PBS-Tween for 1 hour, incubated overnight at 4°C with primary antibodies diluted in 5% skim milk in PBS-Tween, washed three times with PBS-T, incubated 1 hour at room temperature with HRP-conjugated secondary antibodies diluted in 5% skim milk in PBS-Tween, and washed three times with PBS-T. Finally, proteins were detected using enhanced chemiluminescence (Super Signal West Pico Plus, Thermo Scientist, # 34580) and chemiluminescence was imaged on a G:Box chemi XX6 (Syngene).

Antibodies used: BRCA1 (Cell Signaling, #9010T, diluted at 1:1000), PALB2 (generous gift from Jean-Yves Masson), Myc (Cell Signaling, #2276, diluted at 1:1000), TRF2 (Novus, #NB110-57130, diluted at 1:1000), Ku80 (Cell Signaling, #2753, diluted at 1:1000), ß-actin (Cell Signaling, #8457, diluted at 1:5000), secondary anti-Mouse-HRP (Cell Signaling, #70765, diluted at 1:3000), and secondary anti-Rabbit-HRP (Cell Signaling, #7074P2, diluted at 1:3000).

#### Immunofluorescence

For immunofluorescence, 15,000 HT1080, HT1080-BRCA1^KO^, or HT1080-PALB2^KO^ cells were plated on coverslips 24 hours prior to treatment. Cells were either irradiated with 8Gy of ionizing radiation or left unirradiated. Cells were allowed to recover for 2 hours, fixed with 10% formalin for 15 mins, washed with 1x PBS, permeabilized with 0.5% Triton-X and 0.2M HCl for 10 minutes, washed again with 1x PBS and blocked with a 4:1 ratio of 10% BSA in PBS-Tween 0.05% and 20% FBS in PBS for 75 minutes. After blocking, cells were rinsed with PBS and incubated with primary antibodies against Rad51 (Mouse Anti-Rad51 monoclonal, Abcam ab213, diluted at 1:500) or γH2AX (Rabbit mAb Phospho-H2A.X S139, Cell Signaling #9718, diluted at 1:1000) overnight at 4°C. The following day the cells were washed with PBS and incubated with secondary AlexaFluors (Goat anti-Mouse IgG (H+L) Secondary Antibody AlexaFluor 568; Goat anti-Rabbit IgG (H+L) Secondary Antibody AlexaFluor 488, diluted at 1:500) in blocking buffer for 1 hour. Cells were again rinsed with PBS and subsequently dH2O. Coverslips were gently dried and fixed onto slides with ProLongGold + DAPI dye and imaged on an ECHO Revolve Light Microscope. Images were compiled and analyzed using ImageJ.

## QUANTIFICATION AND STATISTICAL ANALYSIS

The data from biological triplicate experiments were expressed as mean ± SDs and analyzed by ANOVA for bar graphs or Mann-Whitney test for scatter plots. Sample size was indicated in corresponding Figure Legends. P < 0.05 was statistically significant (*P < 0.05; **P < 0.01; NS, not significant).

## KEY RESOURCES TABLES

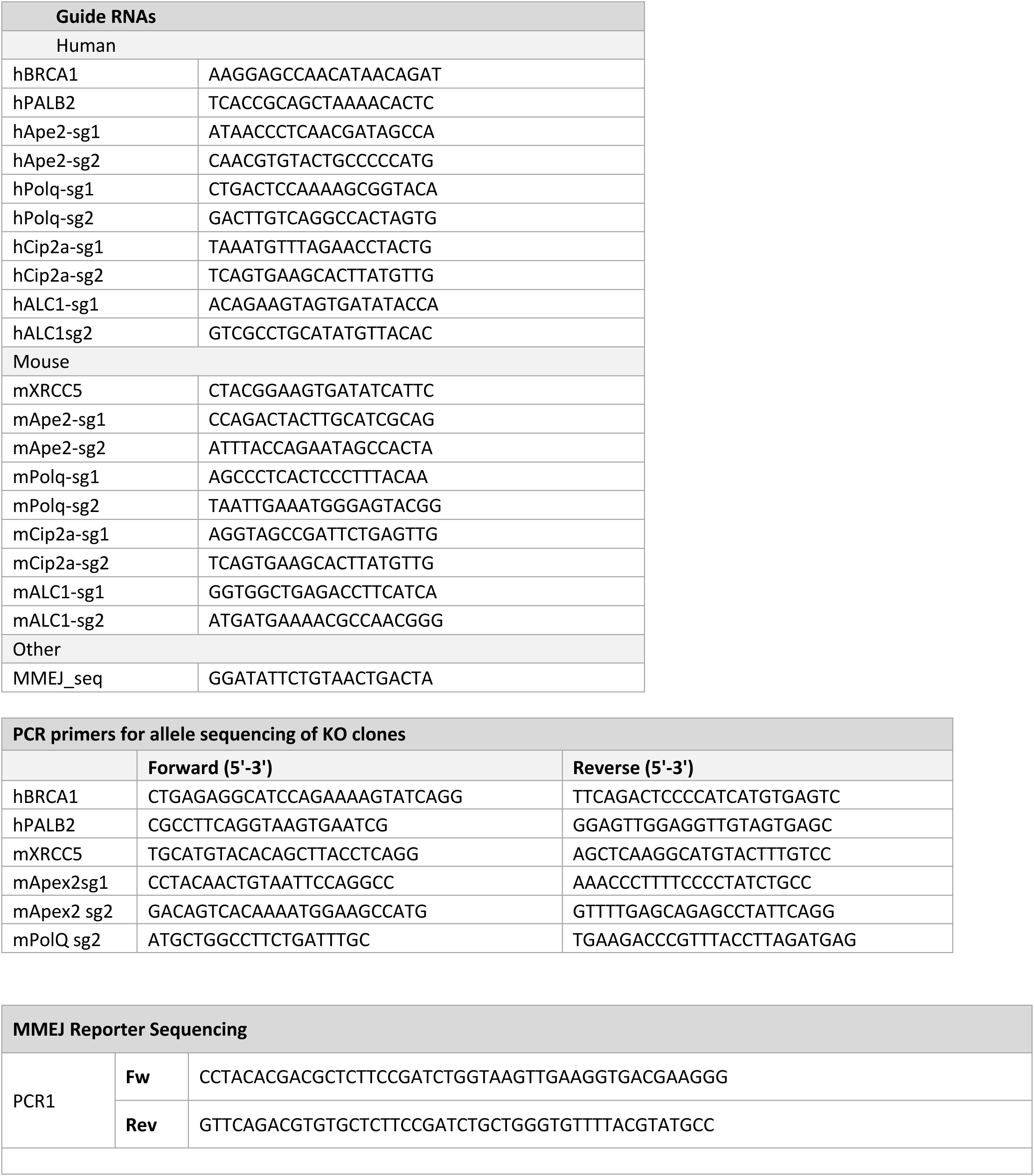
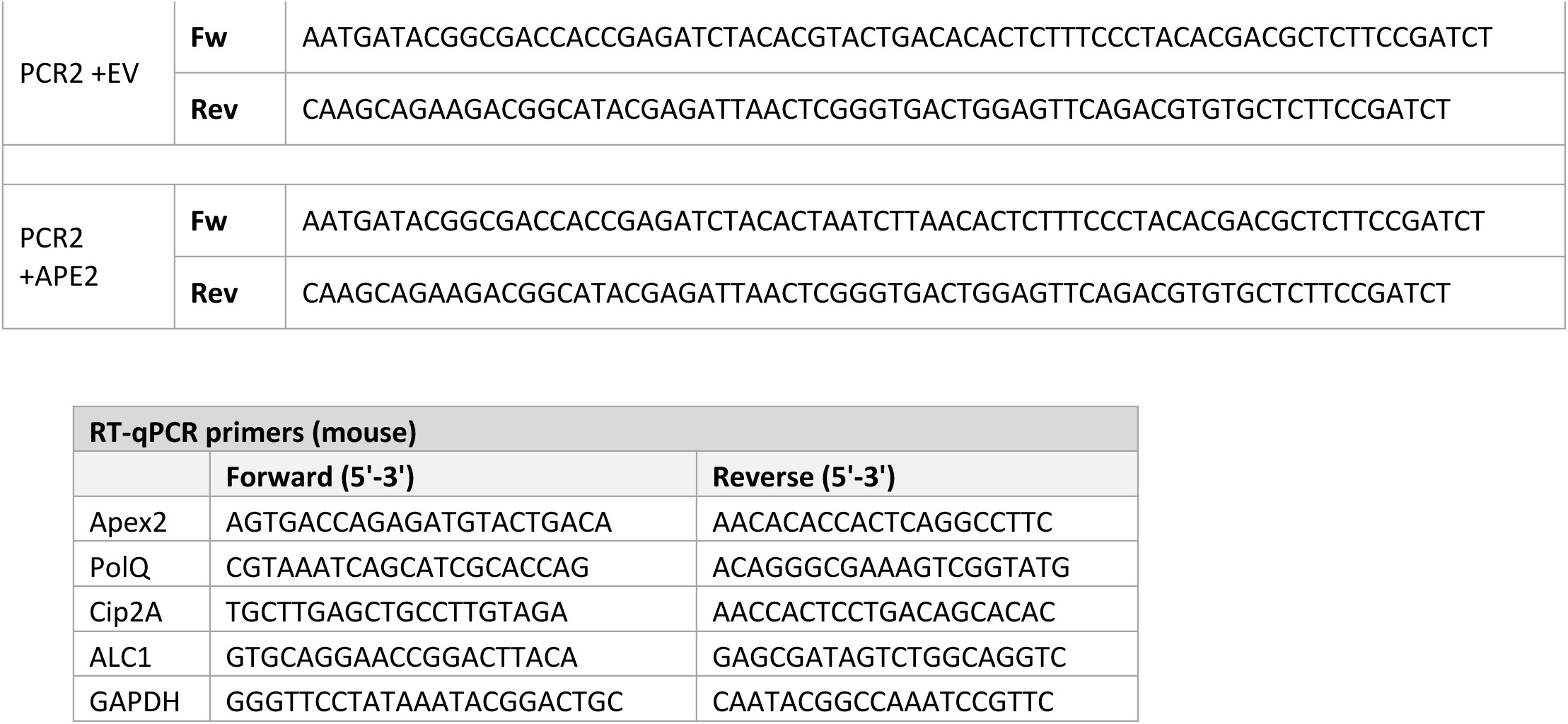

## Acknowledgements

N.A. is supported by the NIH (R35GM143108, R01CA266100), the Boettcher Webb-Waring Foundation, the V Foundation for Cancer Research, and the Glenn Foundation and American Federation for Aging Research. E.T. is supported by the NIH (T32GM142607) and an NSF GRFP fellowship (DGE2040434).

B.N. is supported by an NIH training grant (T32GM008759 and T32GM142607). S.J.B. and R.A. are supported by the Francis Crick Institute, which receives its core funding from Cancer Research UK (FC0010048), the UK Medical Research Council (FC0010048) and the Wellcome Trust (FC0010048).

S.J.B is also funded by a European Research Council (ERC) Advanced Investigator Grant (TelMetab); and Wellcome Trust Senior Investigator and Collaborative Grants. W.L is supported by the NIH (R35GM137798, R01CA244261), American Cancer Society (RSG-20-131-01-DMC, TLC-21-164-01-TLC) and Arkansas Breast Cancer Research Program (AWD00054499 and AWD00053730).

## Contributions

H.F. and N.A. developed the concept for the paper and designed the bulk of the experiments. H.F. and M.K.M. performed and H.F. analyzed the bulk of the experiments of the paper. C.M.S. and J.W.L. designed, performed, and analyzed the micro-irradiation experiments. R.A. and S.J.B. designed, performed, and analyzed the biochemistry experiments. C.S. performed the bioinformatics analyses. B.N. and N.A. performed the CRISPR-Cas9 screens. E.T., B.D., D.D., and R.O. performed experiments and/or made cell lines or plasmids. N.A. and H.F. wrote the manuscript with input from all the other authors.

