## Supplementary Figures for "The APE2 nuclease is essential for DNA double strand break repair by microhomology-mediated end-joining"

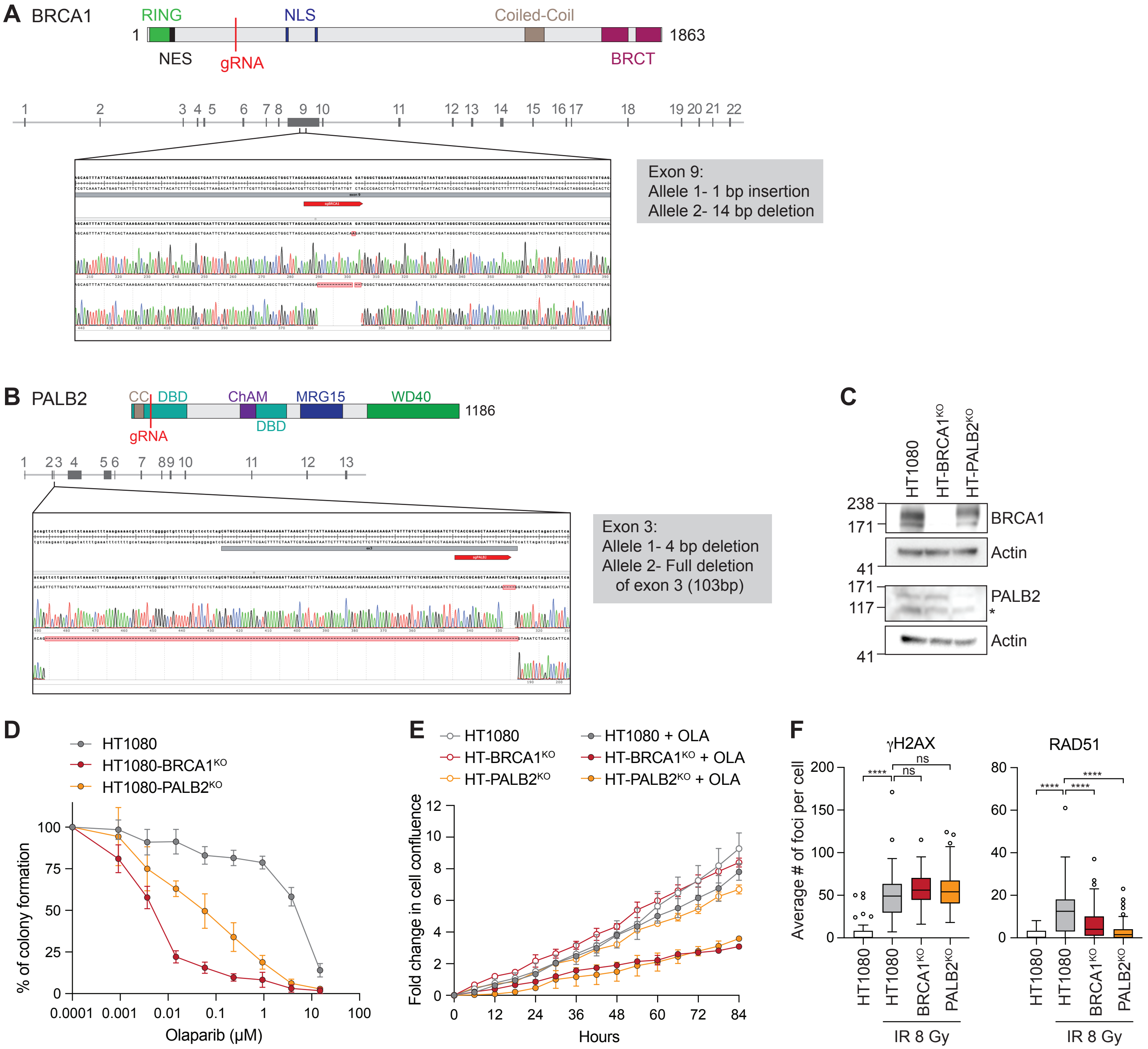


**Figure S1. Creation of isogenic HT1080 and HT1080-BRCA1 and PALB2 knock out clones.**
**A.** Schematic of BRCA1 with location of sgRNA indicated. Representative sequencing data of the 2 mutated alleles of *BRCA1.* **B.** Schematic of PALB2 with location of sgRNA indicated. Representative sequencing data of the 2 mutated alleles of *PALB2*. **C.** Western blot of BRCA1, PALB2 and Actin in parental HT1080 and indicated KO clones. *: unspecific band. **D.** Clonogenic assays of indicated cells treated with increasing concentrations of Olaparib, normalized to no drug. 2 biological replicates, 2 experimental replicates each. Data are mean ± s.d. **E.** Growth curves of indicated cells treated with 1.9μM Olaparib, measured by IncuCyte live microscopy, normalized to 0 hour. 3 independent experiments. Data are mean ± s.e.m. **F.** Quantification of immunofluorescence γH2A.X and RAD51 foci in indicated cells upon irradiation (8Gy). Statistical analysis: One-way ANOVA. ****p<0.0001.

**A**

| | BRCA1 KO | | PALB2 KO | |
|---|---|---|---|---|
| | Log Fold Count | MAGeCK score | Log Fold Count | MAGeCK score |
| XRCC1 | -4.8609 | 2.16E-07 | -5.5297 | 1.98E-08 |
| APEX2 | -4.995 | 2.72E-07 | -4.0405 | 9.82E-08 |
| PARP2 | -0.76862 | 0.016139 | -3.3076 | 3.28E-08 |
| POLQ | -1.4461 | 0.00056255 | -3.1901 | 1.26E-08 |
| RHNO1 | -0.22114 | 0.046974 | -5.3931 | 2.91E-07 |
| ALC1 | -3.531 | 7.45E-05 | -3.4083 | 5.50E-07 |
| PARP1 | -7.2137 | 5.20E-09 | -4.2614 | 2.67E-06 |
| RNASEH2C | -1.2429 | 0.00092039 | -2.5239 | 7.21E-06 |
| PXMP4 | -0.74711 | 0.029079 | -1.4195 | 8.02E-06 |
| PRMT9 | 0.10768 | 0.099055 | -1.0205 | 1.12E-05 |
| NBN | -3.4856 | 1.68E-06 | -1.8518 | 1.44E-05 |
| CIP2A | -4.2451 | 1.66E-05 | -4.0021 | 2.07E-05 |
| MED18 | -0.84742 | 0.022015 | -2.3651 | 2.40E-05 |
| CTC1 | -0.90369 | 0.16279 | -4.3517 | 2.53E-05 |
| SUSD6 | -0.8592 | 0.049793 | -1.6675 | 4.24E-05 |
| LOC1001304 | -0.027574 | 0.52546 | -1.2139 | 4.75E-05 |
| HLA-E | 0.54286 | 0.96536 | -1.4136 | 5.40E-05 |
| LIG3 | -1.9727 | 0.00016037 | -1.8882 | 6.54E-05 |
| PCYT2 | -0.42814 | 0.16748 | -1.5106 | 7.52E-05 |
| TTC4 | -3.7587 | 0.0072999 | -3.177 | 7.80E-05 |
| CENPX | -5.5651 | 1.50E-11 | -0.075795 | 0.49576 |
| UBE2T | -2.4486 | 1.04E-09 | 0.76301 | 0.65891 |
| FAAP24 | -6.1975 | 2.38E-09 | -0.20185 | 0.46912 |
| FANCA | -4.3573 | 4.82E-08 | -0.1678 | 0.23258 |
| APITD1 | -6.7061 | 1.09E-06 | 0.39829 | 0.51807 |
| FANCB | -4.1736 | 4.00E-06 | 0.11388 | 0.64126 |
| SETD2 | -3.2233 | 4.50E-06 | -0.36235 | 0.22616 |
| RAD18 | -1.9932 | 5.04E-06 | 0.018365 | 0.79729 |
| RUNX1 | -1.2304 | 5.98E-06 | -0.93182 | 0.0036222 |
| FANCL | -4.0715 | 6.70E-06 | 0.19127 | 0.87176 |
| PMS2 | -1.3724 | 2.00E-05 | 0.40521 | 0.83124 |
| CHTOP | -1.9392 | 2.78E-05 | -1.0519 | 0.0013315 |
| FAAP100 | -5.2649 | 3.19E-05 | 0.52715 | 0.65421 |
| CENPO | -2.3432 | 3.50E-05 | -0.038538 | 0.32714 |

**B** **BRCA1$^{KO}$ depleted top 20**

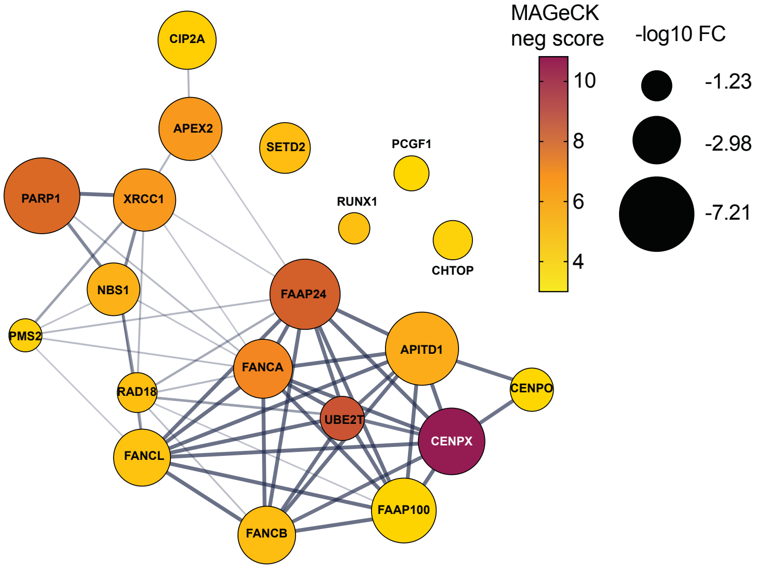


**C** **PALB2$^{KO}$ depleted top 20**

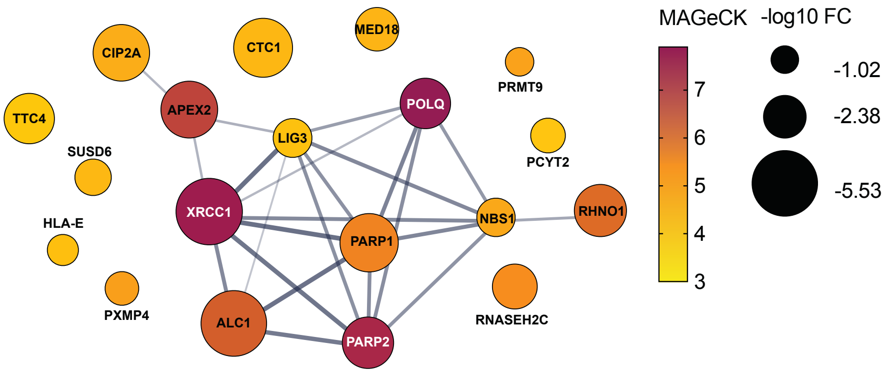


**Figure S2. Genetic interactions of BRCA1 and PALB2.**
**A.** Log fold count and MAGeCK scores in BRCA1 and PALB2 screens for the top 20 candidates of each screen, colored according to LFC of each screen. **B.** String network of the top 20 depleted genes in the BRCA1 screen. **C.** String network of the top 20 depleted genes in the PALB2 screen (1 significant gene, *LOC100130451*, could not be included in the string network).

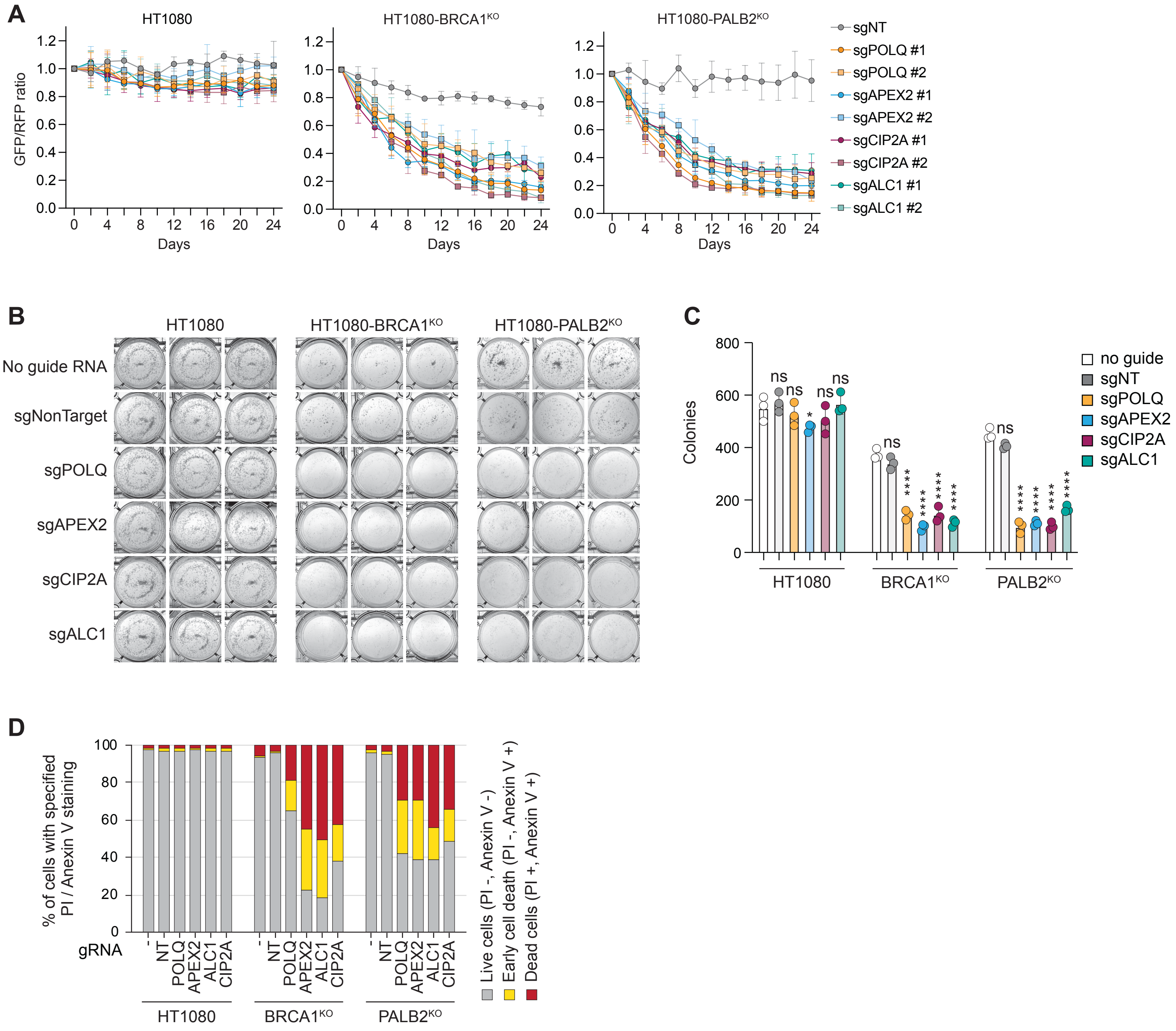


**Figure S3. APEX2, CIP2A and ALC1 are synthetically lethal with HR.**
**A.** Time course result of the growth competition assay. Values are % of RFP+ cells / % of GFP+ cells, normalized to day 0. 3 independent experiments. Data are mean ± s.d. **B-C.** Representative images (B) and quantification (C) of clonogenic assays in HT1080, HT-*BRCA1*KO or HT-*PALB2*KO transduced with iCas9 and the indicated sgRNA and treated with Doxycycline. 3 independent experiments. Data are mean ± s.d. Statistical analysis: One-way ANOVA. *p<0.05, ****p<0.0001. **D.** Quantification of PI /Annexin V staining of HT1080, HT-*BRCA1*KO or HT-*PALB2*KO transduced with iCas9 and the indicated sgRNA and treated with Doxycycline.

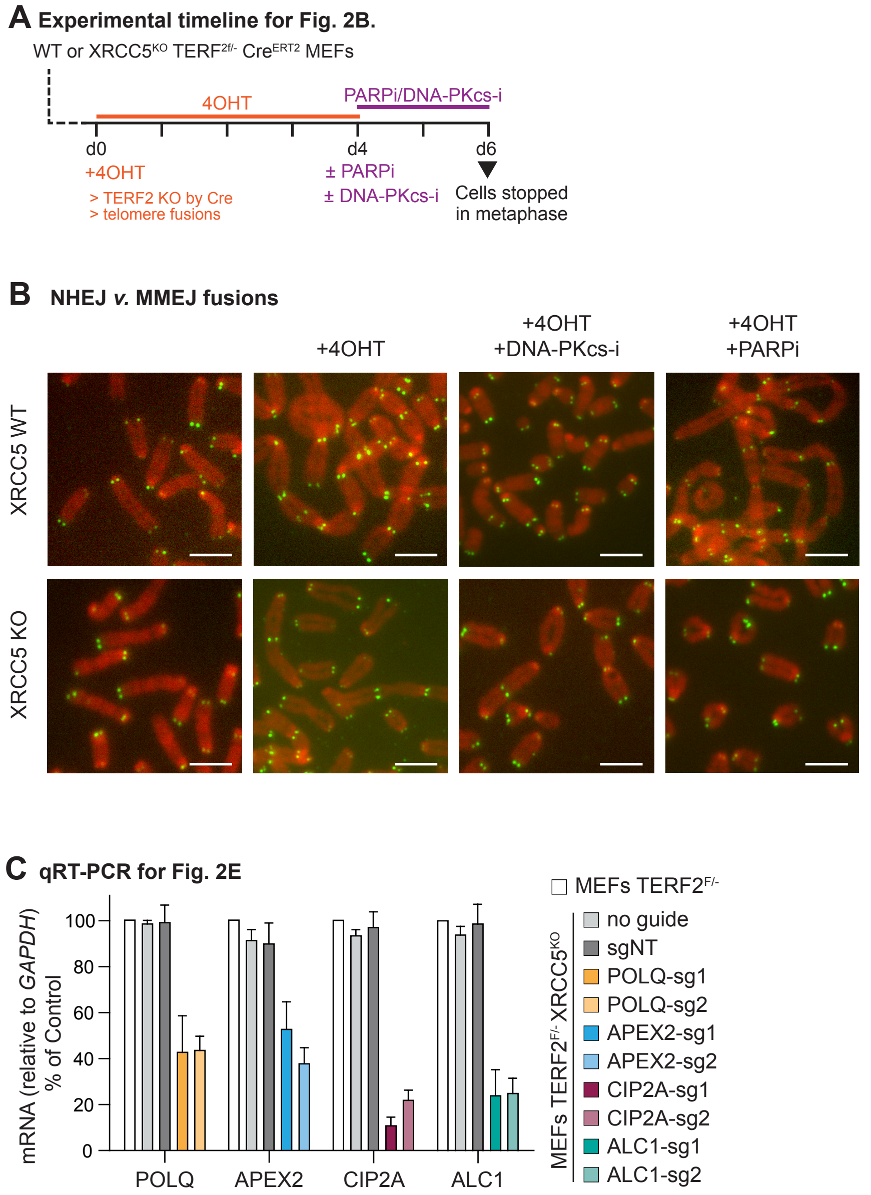


**Figure S4. APE2 is required for MMEJ- but not NHEJ-mediated telomere fusions.**
**A.** Experimental timeline for Figure 2B. **B.** Representative images of metaphases analyzed in Figure 2B. Red: DNA (DAPI), Green: telomere FISH. Scale bar: 5µm. **C.** qRT-PCR quantification of POLQ, APEX2, CIP2A and ALC1, normalized to GAPDH, in MEFs used in Figures 2D and 2E.

**A**

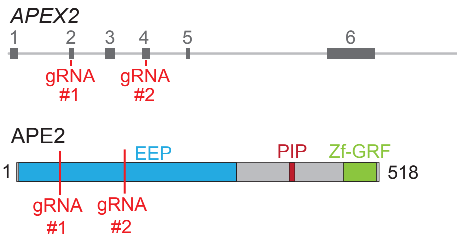


| Clone | Target exon | Allele 1 | Allele 2 |
|---|---|---|---|
| A1.1 | Exon 2 | 1 bp deletion | 4 bp deletion |
| A1.7 | Exon 2 | 1 bp insertion | 1 bp insertion |
| A1.9 | Exon 2 | 4 bp deletion | 1 bp deletion |
| A2.4 | Exon 4 | 1 bp insertion | 1 bp insertion |

**B**

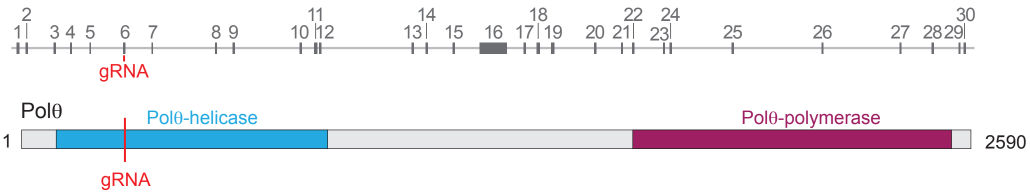


| Clone | Target exon | Allele 1 | Allele 2 |
|---|---|---|---|
| P2.4 | Exon 6 | 1 bp insertion | 17 bp deletion |
| P2.5 | Exon 6 | 1 bp deletion | 1 bp insertion |
| P2.9 | Exon 6 | 5 bp deletion | 1 bp insertion |

**C**

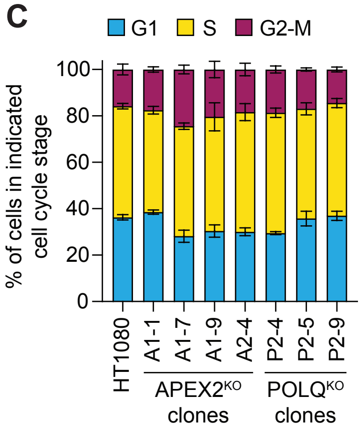


**Figure S5. Characterization of APEX2 and POLQ knock-out clones.**
**A.** Schematic of *APEX2* gene and APE2 protein with location of the sgRNAs #1 and #2 indicated. Representative sequencing data of the 4 different clones. Clones A1.1, A1.7 and A1.9 were obtained with sg*APEX2* #1, clone A2.4 was obtained with sg*APEX2* #2. **B.** Schematic of *POLQ* gene and Polθ protein with location of the sgRNA indicated. **C.** Cell cycle distribution of the different clones. Cells were treated 15 minutes with BrdU, fixed, labelled with Propidium Iodide and anti-BrdU. Cell cycle was determined by flow cytometry analysis of PI and BrdU profiles.

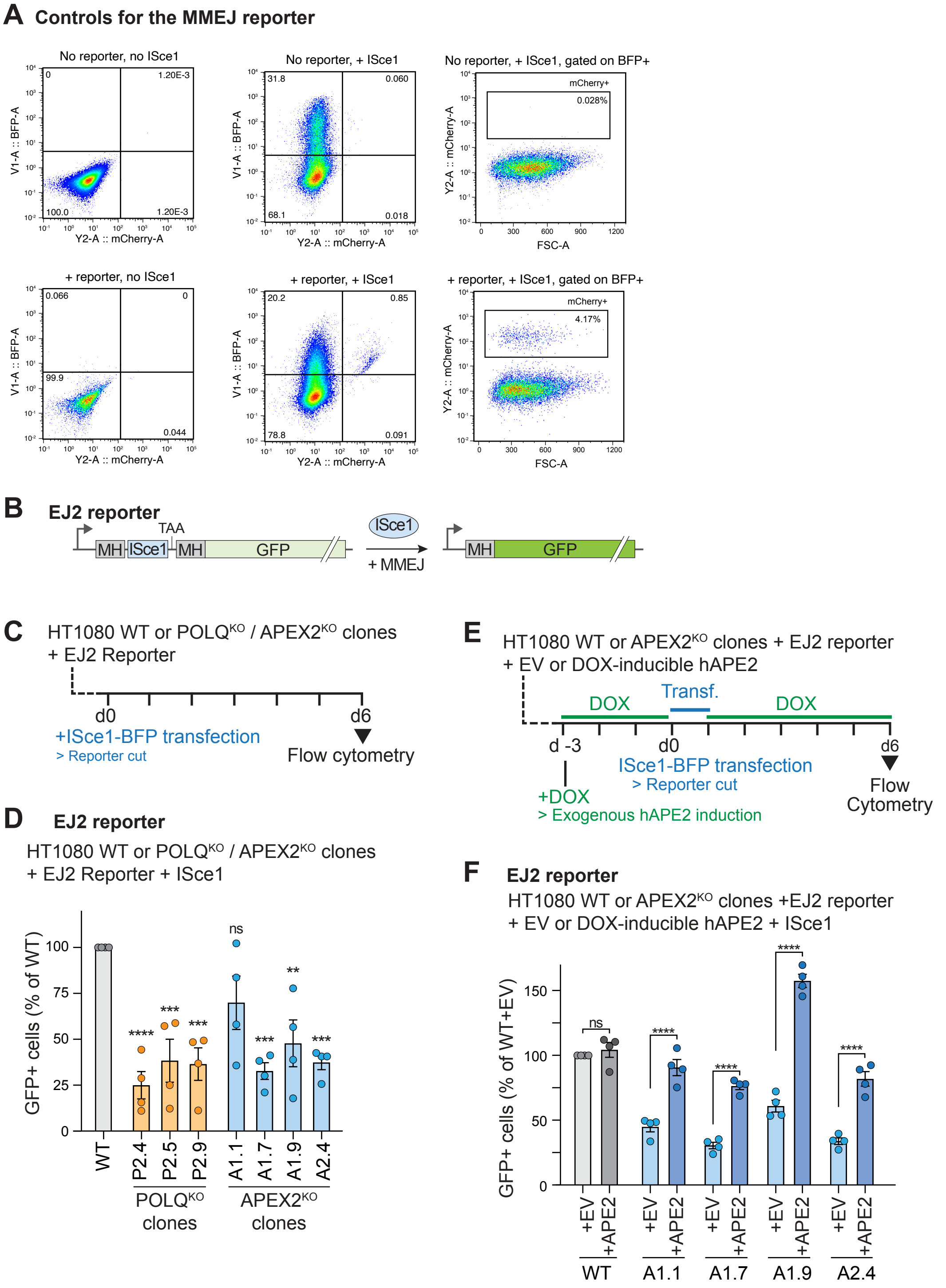


**Figure S6. APE2 is required for MMEJ repair of intra-chromosomal DSBs.**

**A.** Representative flow cytometry analyses controlling the MMEJ reporter. **B.** Schematic of the EJ2 reporter. **C.** Experimental timeline for D. **D.** EJ2 reporter quantification in parental HT1080 and indicated KO clones. GFP+ cells (MMEJ+) were scored in the BFP+ population (I-Sce1+). Values are normalized to WT. 3 independent experiments. Data are mean ± s.e.m. **E.** Experimental timeline for F. **F.** EJ2 reporter quantification in parental HT1080 and *APEX2*$^{KO}$ clones complemented with empty vector (EV) or hAPE2. 4 independent experiments. Data are mean ± s.e.m. Statistical analysis for D and F: One-way ANOVA. **$p<0.01$, ***$p<0.001$, ****$p<0.0001$.

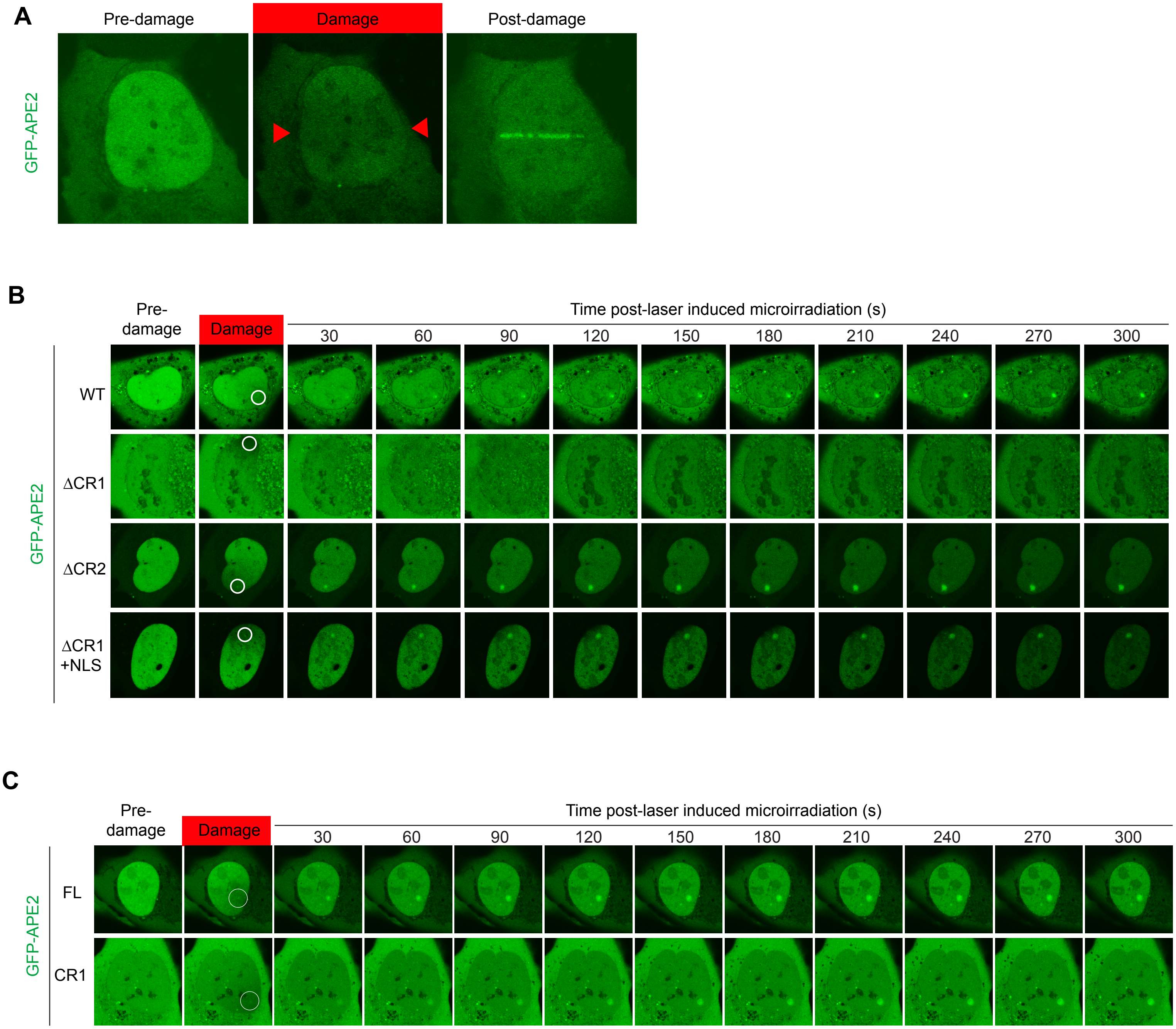


**Figure S7. Recruitment of APE2 to sites of laser-irradiation damage.**
**A.** Recruitment of GFP-APE2 to laser micro-irradiation damage (405nm, 60% power) upon treatment with BrdU (10μM, 20h) in U2OS cells. **B-C.** Time course of recruitment of WT and mutant APE2, as well as the CR1 motif, to laser micro-irradiation sites.
